# SARS-CoV-2 Viral Genes Compromise Survival and Functions of Human Pluripotent Stem Cell-derived Cardiomyocytes via Reducing Cellular ATP Level

**DOI:** 10.1101/2022.01.20.477147

**Authors:** Juli Liu, Yucheng Zhang, Shiyong Wu, Lei Han, Cheng Wang, Sheng Liu, Ed Simpson, Ying Liu, Yue Wang, Weinian Shou, Yunlong Liu, Michael Rubart-von der Lohe, Jun Wan, Lei Yang

## Abstract

Cardiac manifestations are commonly observed in COVID-19 patients and prominently contributed to overall mortality. Human myocardium could be infected by SARS-CoV-2, and human pluripotent stem cell-derived cardiomyocytes (hPSC-CMs) are susceptible to SARS-CoV-2 infection. However, molecular mechanisms of SARS-CoV-2 gene-induced injury and dysfunction of human CMs remain elusive. Here, we find overexpression of three SARS-CoV-2 coding genes, Nsp6, Nsp8 and M, could globally compromise transcriptome of hPSC-CMs. Integrated transcriptomic analyses of hPSC-CMs infected by SARS-CoV-2 with hPSC-CMs of Nsp6, Nsp8 or M overexpression identified concordantly activated genes enriched into apoptosis and immune/inflammation responses, whereas reduced genes related to heart contraction and functions. Further, Nsp6, Nsp8 or M overexpression induce prominent apoptosis and electrical dysfunctions of hPSC-CMs. Global interactome analysis find Nsp6, Nsp8 and M all interact with ATPase subunits, leading to significantly reduced cellular ATP level of hPSC-CMs. Finally, we find two FDA-approved drugs, ivermectin and meclizine, could enhance the ATP level, and ameliorate cell death and dysfunctions of hPSC-CMs overexpressing Nsp6, Nsp8 or M. Overall, we uncover the global detrimental impacts of SARS-CoV-2 genes Nsp6, Nsp8 and M on the whole transcriptome and interactome of hPSC-CMs, define the crucial role of ATP level reduced by SARS-CoV-2 genes in CM death and functional abnormalities, and explore the potentially pharmaceutical approaches to ameliorate SARS-CoV-2 genes-induced CM injury and abnormalities.

## Introduction

Coronavirus disease 2019 (COVID-19), which is caused by the severe acute respiratory syndrome coronavirus 2 (SARS-CoV-2), has unprecedentedly affected health and economy world widely (1, 2). Since the end of 2019, SARS-CoV-2 has infected over 339 million people in the world, and in the US, SARS-CoV-2 has caused over 850,000 deaths. Although the primary cause of death by SARS-CoV-2 infection is respiratory failure, cardiac complications, including acute myocardial injury, myocarditis, arrhythmias and even sudden death, prominently contributed to the overall SARS-CoV-2-caused mortality (3–5). Concomitant cardiovascular disorders have been observed in 8-25% of overall SARS-CoV-2 infected population, and particularly with a high frequency among dead patients (6). Additionally, an increased mortality rate was observed in patients with pre-existing cardiovascular disorders (7, 8). Recent studies documented the vulnerability of human myocardium infection by SARS-CoV-2, confirming that SARS-CoV-2 directly infects CMs in COVID-19 patients with myocarditis (9). SARS-CoV-2 virus infects and enters human cells through binding to three membrane proteins, including the Angiotensin Converting Enzyme 2 (ACE2), Transmembrane Serine Protease 2 (TMPRSS2)(10, 11) and Transmembrane Serine Protease 4 (TMPRSS4)(12). The high affinity of SARS-CoV-2 spike (S) protein with ACE2 makes human organs/tissues with high level of ACE2 expression as the primary targets attacked by SARS-CoV-2, such as the lung, small intestine, testis, kidney and heart (13). Recently, human pluripotent stem cell-derived cardiomyocytes (hPSC-CMs) were naturally utilized as an *in vitro* platform to evaluate the pathological effects of SARS-CoV-2 on human heart muscle cells (14). SARS-CoV-2 infects human CMs through ACE2. SARS-CoV-2- infected hPSC-CMs exhibited prominently increased cell death (15, 16), fractionated sarcomeres (17), abnormal electrical and mechanical functions (18) and inflammation (15), which recapitulated the CM injuries in COVID-19 patients with myocarditis (9). SARS-CoV-2 genome encodes up to 27 genes. Currently, the molecular mechanisms, by which SARS-CoV-2 genes interact with host protein networks to influence survival and functions of human CMs, still largely remain elusive.

In this study, three SARS-CoV-2 viral coding genes, Nsp6, Nsp8 and M, were overexpressed in hPSCs (human ESCs and iPSCs) derived CMs. The global impacts of SARS-CoV-2 viral genes on the transcriptome of hPSC-CMs were determined by whole mRNA-seq. We found Nsp6, Nsp8 or M overexpression in hPSC-CMs and SARS-CoV-2 viral infection concordantly affected the transcriptome of hPSC-CMs, with activated cellular injury and immune signaling and reduced cardiac function pathways. Nsp6, Nsp8 or M overexpression in hPSC-CMs prominently increased apoptosis, and compromised calcium handling and electrical properties. Genome-wide interactome analysis found Nsp6, Nsp8 and M could all interact with proteins of the host hPSC-CMs, particularly ATPase subunits, which led to significantly reduced ATP level. Given that reduced ATP level could impair intracellular Ca^2+^ signaling and contractility of CMs, we then tested pharmaceutical strategies to enhance the cellular ATP levels of hPSC-CMs overexpressing Nsp6, Nsp8 or M. Two FDA-approved drugs, ivermectin and meclizine, significantly reduced cell death and dysfunctions of hPSC-CMs with overexpression of SARS-CoV-2 genes. Overall, SARS-CoV-2 genes Nsp6, Nsp8 and M exhibited detrimental effects on the whole transcriptome and interactome of hPSC-CMs, and the reduced cellular ATP level plays a key role in SARS-CoV-2 genes-induced CM death and abnormalities.

## Methods

A detailed methods section is in the Supplemental Appendix.

### Human Pluripotent stem cell culture and differentiation

Human embryonic stem cell (hESC) line H9 and human induced pluripotent stem cell (hiPSC) line S3 were cultured on Matrigel (BD Biosciences)-coated plates in mTesR1 medium (19, 20). For cardiomyocyte differentiation, cells were differentiated by following a published protocol (21) with minor modification.

### Lentivirus production and cell transduction

The lentiviral vectors pLVX-EF1alpha-IRES-Puro-2xStreg-SARS-CoV-2 (Nsp6, Nsp8, M) (Addgene plasmids #141395, #141372, #141374) were transfected into the HEK293T cells with packaging plasmids. After 48 hrs, viral supernatant was collected. H9 hESCs or S3 hiPSCs cultured in mTesR1 medium were incubated with viral media for 4 hrs. Same infection was repeated after 24 hrs. Puromycin was added to select puromycin-resistant H9 hESCs and S3 hiPSCs clones after 48 hrs of virus infection. Empty vector infected hPSCs were used as control.

### Whole mRNA-seq

Total RNA was evaluated for quantity and quality by using Agilent Bioanalyzer 2100. The pooled cDNA libraries were sequenced by a NovaSeq 6000 sequencer at 300pM final concentration for 100b paired-end sequencing (Illumina, Inc.). Approximately 30-40M reads per library were generated. A Phred quality score (Q score) was used to measure the quality of sequencing. More than 90% of the sequencing reads reached Q30 (99.9% base call accuracy).

### Co-immunoprecipitation mass spectrometry (Co-IP-MS)

Total cell proteins were extracted by using Pierce™ Classic Magnetic IP/Co-IP Kit (Thermo Scientific, 88804, USA) according to the manuals. Protein Co-immunoprecipitation (Co-IP) was performed by using Pierce™ Classic Magnetic IP/Co-IP Kit as well. Co-IP protein samples analysis were performed by using Western blotting or submitted to the Proteomics Core Facility at the Indiana University School of Medicine (IUSM) for mass spectrometry analysis.

### Quantification and Statistical Analysis

Data comparison between two groups (gene overexpression versus control) was conducted using an unpaired two-tailed *t*-test. All data were presented as mean ± S.D. from at least three independent experiments. Differences with *P* values less than 0.05 were considered significant.

## Results

### Prediction of SARS-CoV-2 coding genes that potentially affect functions of human cardiomyocytes (CMs)

Myocardial injury and dysfunction are observed in patients infected by the SARS-CoV-2 virus(22). The 30kb SARS-CoV-2 viral genome encodes up to 14 open-reading frames (Figure 1A). Currently, the host responsive mechanisms of human heart muscle cells to SARS-CoV-2 viral genes still remain elusive. To predict the impact and mechanisms of SARS-CoV-2 coding genes on human cardiac injuries and dysfunctions, we analyzed the published datasets from a previous study(23), which comprehensively defined the interactive map of all SARS-CoV-2 proteins in human HEK-293T/17 cells. Here, we only focus on the SARS-CoV-2 genes, which mainly interact with proteins that are highly expressed in human CMs. By overlapping the 332 high-confident SARS-CoV-2 interactors in the HEK-293T/17 cells with the 2198 proteins that are highly expressed in the human cardiomyocytes (human protein atlas, http://www.proteinatlas.org), we found three SARS-CoV-2 coding genes, Nsp6, M and Nsp8, could selectively interact with the proteins essential for survival and function of human CMs (Figure 1B). Therefore, we decided to pursue the functional studies of those three SARS-CoV-2 genes in hPSC-CMs. Firstly, three human ES and iPS cell lines overexpressing M, Nsp8 and Nsp6, respectively, were established by using lentivirus, followed with differentiation into contracting CMs for further assessments (Figure 1C). This strategy allows uniformly high gene expression in hPSC-CMs compared with the AAV9-mediated gene delivery into differentiated hPSC-CMs. Since SARS-CoV-2 virus infects human cells through binding with ACE2, TMPRSS2(10, 11) or TMPRSS4(12), we profiled their expression dynamics during CM differentiation from WT hESCs using qRT-PCR (Figure 1D). Cardiac troponin T (CTNT) is a marker of CMs (Figure 1D). ACE2 was lowly expressed in hESCs, but rapidly increased in hESCs-derived CMs (hESC-CMs). TMPRSS2 expression level dropped during CM differentiation (Figure 1D), whereas TMPRSS4 expression exhibited a fluctuant patterning (Figure 1D). Our single cell RNA-seq data from hESC-CMs (24) confirmed the relatively higher expression level of ACE2 than those of TMPRSS2 and TMPRSS4 in hESC-CMs (Figure 1E). Similar as hESC-CMs, adult human heart tissues express higher level of ACE2 than TMPRSS2 and TMPRSS4 (Supplementary Figures 1A-C). Furthermore, pseudotyped SARS-CoV-2 virus could infect WT hESC-CMs (Figure 1F), which was consistent with recent reports of direct SARS-CoV-2 infection of hPSC-CMs (15, 16).

**Figure 1.**
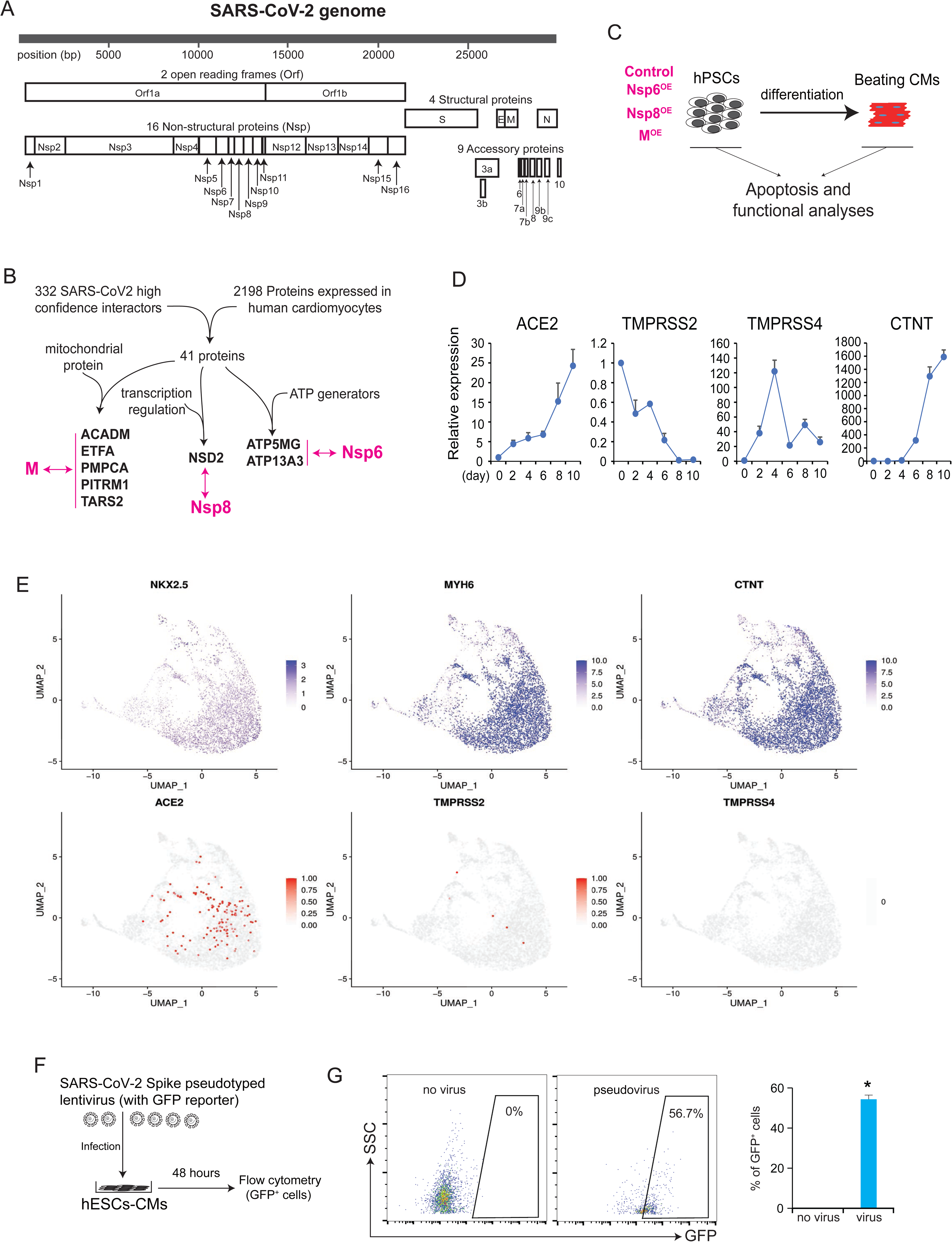
Prediction of SARS-CoV-2 viral genes affecting human cardiomyocytes and confirmation of infection by SARS-CoV-2 spike pseudotyped lentivirus. **(A)** Coding genes annotation of the SARS-CoV-2 genome. **(B)** Strategies to predict SARS-CoV-2 genes potentially affecting functions of human cardiomyocytes. The double-headed arrows indicate the potential interactions between SARS-CoV-2 proteins and essential proteins in human cardiomyocytes. **(C)** Stable hPSC cell lines were established to study the functions of SARS-CoV-2 genes in human CMs. OE, overexpression. **(D)** RT-qPCR detection of SARS-CoV-2 virus receptor genes ACE2, TMPRSS2, TMPRSS4 and the cardiomyocyte marker CTNT during cardiac differentiation from human ESCs. RNA samples were collected every 2 days from day 0 to day 10 of differentiation. Beating cardiomyocytes were observed at day 7. All dots are shown as mean ± SD. (n=3). **(E)** Gene expression profiles of SARS-CoV-2 viral receptor genes ACE2, TMPRSS2/3, and cardiomyocyte markers NKX2-5, MYH6, CTNT in hESC-derived cardiomyocytes based on our scRNA-seq data^33^. **(F)** Flow cytometry analysis of GFP^+^ cells in hESC-CMs after infection with SARS-CoV-2 Spike pseudotyped lentivirus with GFP reporter. After 48 hours of infection, flow cytometry was performed to detect GFP^+^ cells. Cells without virus infection were the blank control cells. All bars are shown as mean ± SD. (n=3). A two-tailed unpaired *t*-test was used to calculate *P*-values: \**P* < 0.05 (vs. no virus).

### Whole mRNA-seq reveals the global impact of SARS-CoV-2 viral genes on the transcriptome of human cardiomyocytes

Nsp6^OE^, Nsp8^OE^ or M^OE^ and control hESCs (OE, overexpression; Figures 2A, Supplementary Figure 1D) were differentiated into CMs, followed with a published protocol (25) to enrich CMs till over 90% purity by adding lactate medium. Empty lenti-vector infected hPSCs were used as control. Next, whole mRNA-seq was performed to profile differentially expressed genes (Figure 2B). We found Nsp6^OE^, Nsp8^OE^ or M^OE^ hESC-CMs exhibited significantly changed gene expression profile when compared with control CMs (Figures 2C-E, Table S1). Interestingly, gene ontology (GO) analysis showed that Nsp6^OE^, Nsp8^OE^ or M^OE^ globally affect the transcriptome of hESC-CMs, including activated immune response, protein modification/ubiquitination, defense response, regulation of proliferation and extracellular matrix organization etc. in hESC-CMs. (Figures 2F-G, Supplementary Figures 2A-F). Moreover, in hESC-CMs, genes co-upregulated by Nsp6^OE^, Nsp8^OE^ and M^OE^ could be enriched into multiple signaling pathways, including p53 signaling, fibrosis, oxidative stress response, and virus entry via endocytic pathways (Figures 2H-I). These data indicated that Nsp6^OE^, Nsp8^OE^ and M^OE^ could trigger stress-related signaling pathways in human CMs. Additionally, Nsp6^OE^, Nsp8^OE^ and M^OE^ differentially affected the expression of genes associated with senescence, NF-kB and STAT3 signaling pathways (Figure 2J), suggesting the diverse influences of SARS-CoV-2 genes on the transcriptome of host hESC-CMs.

**Figure 2.**
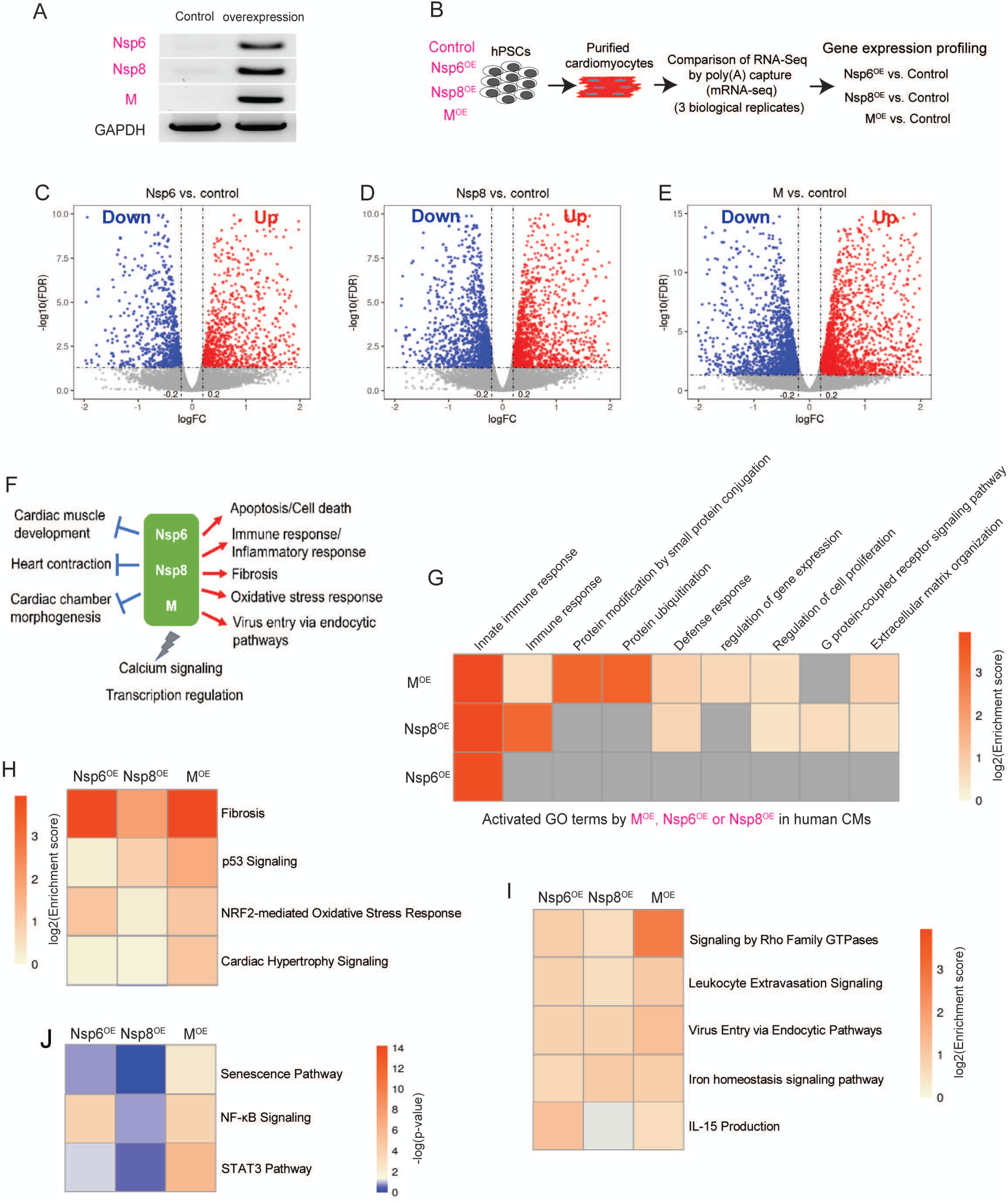
Whole mRNA-seq reveals the impact of SARS-CoV-2 viral genes on the transcriptome of hESC-derived cardiomyocytes. **(A)** RT-PCR detection of SARS-CoV-2 genes Nsp6, Nsp8 and M in the stable overexpression (OE) H9 hESC lines. **(B)** The scheme of whole mRNA-seq to globally study the impact of of SARS-CoV-2 viral genes on purified hESC-CMs. **(C-E)** Volcano blots showing differential expressed genes caused by Nsp6^OE^ **(C)**, Nsp8 ^OE^ **(D)** and M ^OE^ **(E)** in hESC-CMs. **(F)** Global view of transcriptomic changes induced by Nsp6^OE^, Nsp8 ^OE^ and M ^OE^ in hESC-CMs. **(G)** Gene ontology (GO) analyses of the upregulated genes induced by Nsp6^OE^, Nsp8 ^OE^ and M ^OE^ in hESC-CMs. **(H-I)** Canonical signaling pathways analyses of the upregulated genes induced by Nsp6^OE^, Nsp8 ^OE^ and M ^OE^ in hESC-CMs. (**J**) Canonical signaling pathways analyses of the downregulated genes induced by Nsp6^OE^, Nsp8 ^OE^ and M ^OE^ in hESC-CMs.

Next, we compared the transcriptomic alternations due to these fthree individual SARS-CoV-2 proteins with gene expression changes of the hiPSC-CMs infected by the wild type SARS-CoV-2 virus from a previously published mRNA-seq dataset (Sharma *et al.*, 2020) (Figure 3A). We found that both up- and down-regulated DEGs caused by SARS-CoV-2 infection were significantly enriched in DEGs by Nsp6^OE^, Nsp8^OE^ and M^OE^ (Figure 3B-C), which were enriched in apoptosis, inflammation/immune and heart contraction pathways. Together, Nsp6^OE^, Nsp8^OE^ and M^OE^ and SARS-CoV-2 infection globally influenced the transcriptome of hPSC-CMs by activating cellular injury responses, and affecting CM functions, suggest the important role of thee three SARS-CoV-2 genes in SARS-CoV-2 infection-caused CM injuries. Interestingly, ∼70% of DEGs in SARS-CoV-2-infected hiPSC-CMs were not prominent in individual Nsp6^OE^, Nsp8^OE^ and M^OE^ hiPSCs (Figure 3G-H, grey areas), which were enriched into cellular metabolism, regulation of transcription, apoptosis (Figure 3I), and ATP/fatty acid/amino acid metabolic process (Figure 3J). These significant differences at transcriptomic level indicate that other SARS-CoV-2 genes, except Nsp6/Nsp8/M, could also differentially affect the transcriptome of host hiPSC-CMs, likely via targeting different genes or pathways.

**Figure 3.**
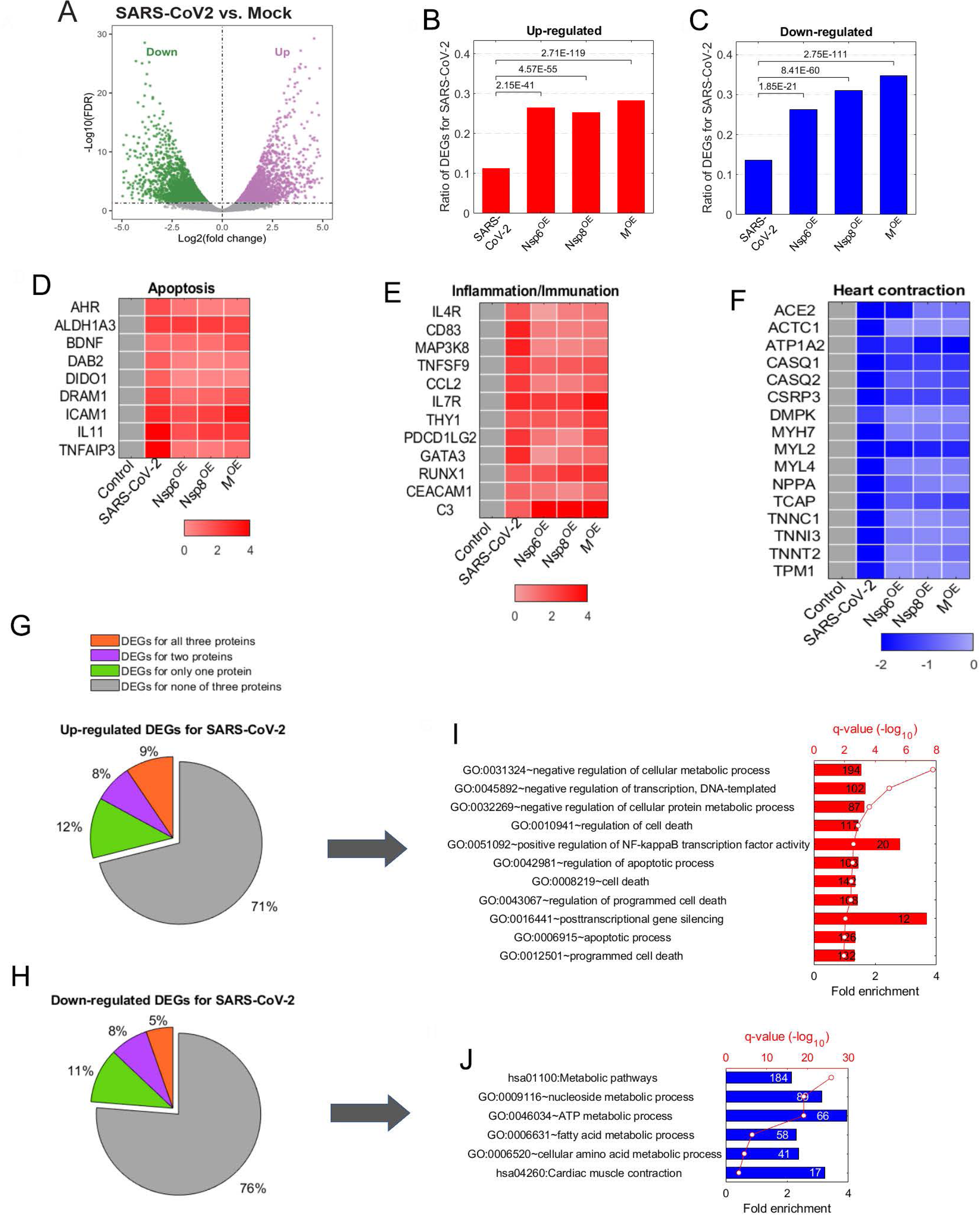
Whole mRNA-seq reveals SARS-CoV-2 viral genes-induced cell death and dysfunctions in hESC-derived cardiomyocytes. **(A)** Volcano blots showing differential expressed genes in hiPSCs (SARS-CoV-2 vs. Mock). (B-C) Comparison of DEGs between hESC-CMs (Nsp6^OE^, Nsp8^OE^ and M^OE^ vs. Control) and hiPSCs (SARS-CoV-2 vs. Mock). The fold enrichment (F.E.) around 2 with remarkable *p*-value indicates the significant overlap between two gene sets. Heat map analyses of apoptotic genes in **(D)**, inflammation genes in **(E)**, and heart contraction genes in **(F).** **(G)** Frequencies of SARS-CoV-2 infection up-regulated DEGs identified in Nsp6^OE^/ Nsp6^OE^/M^OE^ or not. **(H)** Frequencies of SARS-CoV-2 infection down-regulated DEGs identified in Nsp6^OE^/ Nsp6^OE^/M^OE^ or not. **(I)** GO analysis of DEGs solely up-regulated by SARS-CoV-2 infection. **(J)** GO analysis of DEGs solely down-regulated by SARS-CoV-2 infection. All bars are shown as mean ± SD. (n=3). A two-tailed unpaired *t*-test was used to calculate *P*-values: \**P* < 0.05 (vs. Control).

### SARS-CoV-2 genes induce cell death in hPSC-derived cardiomyocytes

The whole mRNA-seq results indicated that overexpression of SARS-CoV-2 viral genes could increase expressions of cell death/apoptosis associated genes in hESC-CMs. Therefore, we next assessed apoptosis of hESC-CMs. Firstly, Nsp6^OE^, Nsp8^OE^, M^OE^ and control hESCs were differentiated into CMs using a 2D monolayer differentiation method(21, 24) (Figure 4A). After 12 days’ CM differentiation, flow cytometry was conducted to detect the ratios of TUNEL^+^ cells in the CTNT^+^ hESC-CMs. We found significantly increased ratios of TUNEL^+^ cells in the Nsp6^OE^, Nsp8^OE^ and M^OE^ hESC-CMs when compared with control hESC-CMs (Figures 4B-C). Immunofluorescent staining on the 2D cultured hESC-CMs also found increased percentages of TUNEL^+^ cells in Nsp6^OE^, Nsp8^OE^ and M^OE^ hESC-CMs compared to that of control hESC-CMs (Figures 4D-E). Since hESC-CMs were dissociated into single cells for flow cytometry analysis, some late-stage apoptotic cells tended to be broken during dissociation, the flow cytometry quantification of apoptotic cells gave rise to a lower percentage value than that by immunostaining. To further confirm the results of TUNEL assay, we quantified the ratios of Annexin V^+^ CMs and observed increased ratios of Annexin V^+^ CMs in Nsp6^OE^, Nsp8^OE^ or M^OE^ hESC-CMs compared to control hESC-CMs (Figures 4F-G). Annexin V has a strong affinity for phosphatidylserine (PS) residues on the surface of the cell that is an early marker of apoptosis. Finally, we quantified the ratios of cleaved-CASP3^+^ CMs (Supplementary Figure 3). Cleaved-CASP3 is an activation form of Caspase-3 responsible for apoptosis execution. Increased percentages of cleaved-CASP3^+^ CMs were found in Nsp6^OE^, Nsp8^OE^ or M^OE^ hESC-CMs when compared with control hESC-CMs, shown by the data from flow cytometry (Supplementary Figure 3A) and immunostaining (Supplementary Figures 3B-C). Together, these results demonstrate that Nsp6^OE^, Nsp8^OE^ and M^OE^ could induce significant cell death of hESC-CMs under the 2D culture condition.

**Figure 4.**
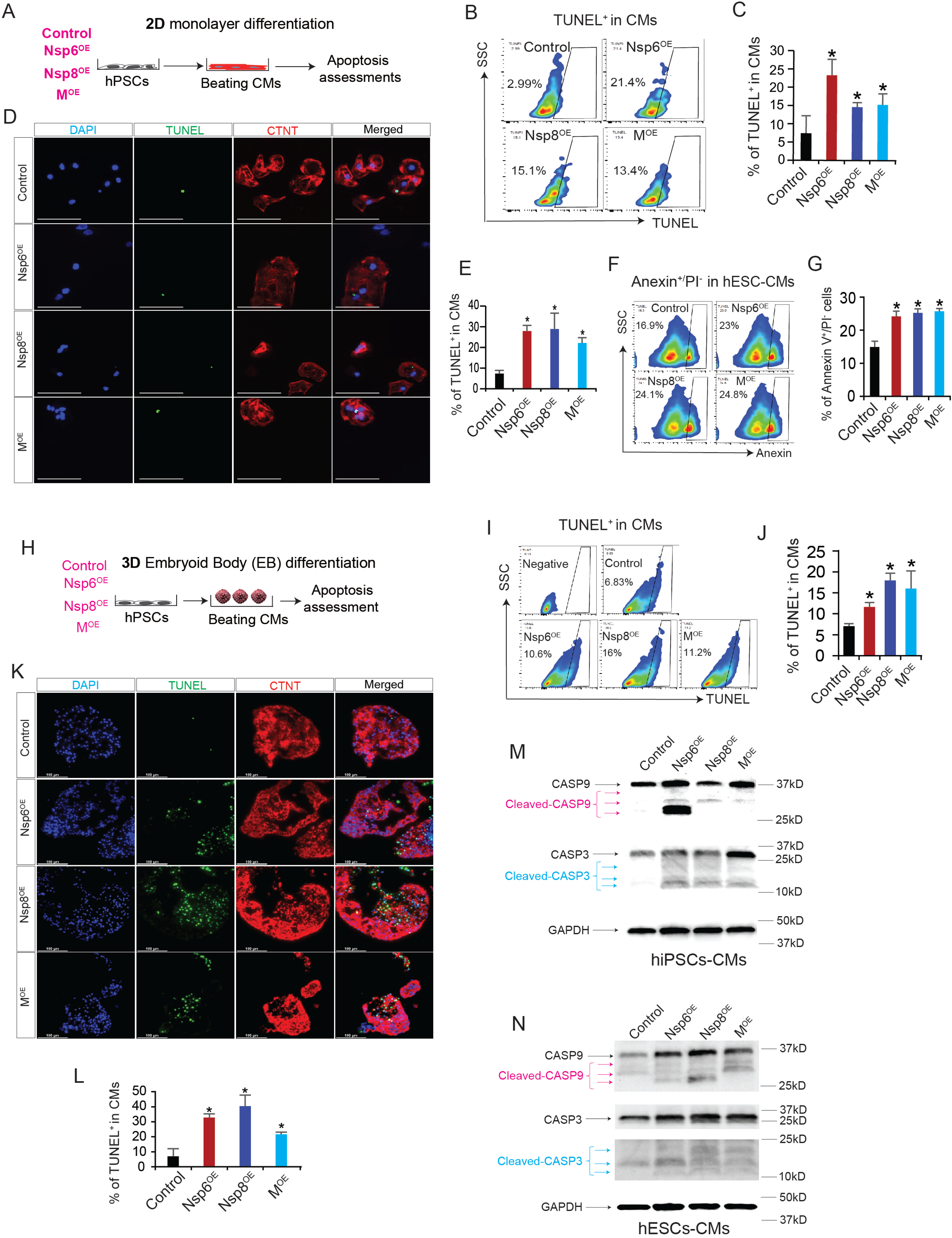
Overexpression of SARS-CoV-2 genes induces cell death of hPSCs- derived cardiomyocytes. **(A)** The scheme cardiomyocytes differentiation from hESCs in 2D monolayer condition, followed with apoptosis assessments. **(B-C)** Flow cytometry analysis of the ratios of TUNEL^+^ cells in hESCs-derived cardiomyocytes (CTNT+). (C) All bars are shown as mean ± SD. (n=4). A two-tailed unpaired *t*-test was used to calculate *P*-values: \**P* < 0.05 (vs. Control). **(D)** Representative immunostaining images of TUNEL^+^ and CTNT^+^ cells in hESCs-derived cardiomyocytes with overexpression of SARS-CoV-2 genes. Scale bar, 100 µm. **(E)** Statistical data analysis of immunostaining results from (**D**). All bars are shown as mean ± SD. (n=3). A two-tailed unpaired *t*-test was used to calculate *P*-values: \**P* < 0.05 (vs. Control). **(F-G)** Flow cytometry analysis of apoptotic cells in hESC-derived cardiomyocytes using Annexin-V-PI double staining. All bars are shown as mean ± SD. (n=3). A two-tailed unpaired *t*-test was used to calculate *P*-values: \**P* < 0.05 (vs. Control). **(H)** The scheme cardiomyocytes differentiation from hiPSCs in 3D Embryoid Body (EB) condition, followed with apoptosis assessments. **(I-J)** Flow cytometry analysis of TUNEL^+^ cells in the CM (CTNT+) population of hiPSCs-derived EBs. All bars are shown as mean ± SD. (n=4). A two-tailed unpaired *t*-test was used to calculate *P*-values: \**P* < 0.05 (vs. Control). **(K)** Representative immunostaining images of TUNEL^+^ cells in hiPSCs-derived EBs containing CTNT^+^ CMs. Scale bar, 100 µm. **(L)** Statistical data analysis of immunostaining results from (**K**). All bars are shown as mean ± SD. (n=3). A two-tailed unpaired *t*-test was used to calculate *P*-values: \**P* < 0.05 (vs. Control). **(M-N)** Western blotting detection of CASP3, cleaved CASP3, CASP9 and cleaved CASP9 protein expression in hiPSC-CMs (**M**) and hESC-CMs (**N**).

Next, in order to recapitulate human myocardium-like tissues, 3D beating embryoid bodies (EBs) were differentiated from hiPSCs by using our established differentiation system (26, 27) (Figure 4H). Prominently increased percentages of TUNEL^+^ CMs were found in Nsp6^OE^, Nsp8^OE^ or M^OE^ hiPSC-derived 3D EBs when compared to control hiPSC-EBs by using flow cytometry and immunofluorescent staining (Figures 4I-L), which were similar with the results from 2D condition-derived CMs. Lastly, the protein levels of apoptotic markers, cleaved-CASP3 and cleaved-CASP9, significantly increased in hiPSC/hESC-EBs with Nsp6^OE^, Nsp8^OE^ or M^OE^ when compared with control EBs (Figures 4M-N). All these results demonstrate that overexpression of SARS-CoV-2 genes can induce cell death of hPSC-CMs.

### Co-IP Mass Spectrometry (MS) reveals the interactive protein networks of hPSC-CMs with SARS-CoV-2 genes

Strep-tagged SARS-CoV-2 genes and GFP^31^ allowed pull down of all interactive proteins in the host cells. We performed Co-IP MS to capture and identify the interactome of Nsp6, Nsp8 or M in hESC-CMs, respectively (Figure 5A). The complete list of all interactors with Nsp6, Nsp8 or M proteins in hESC-CMs could be found in the Supplementary Table S2. Gene Ontology (GO) analyses of the pulled down proteins revealed that Nsp6, Nsp8 and M could all interact with protein factors associated with immune response, viral process, and vesicle-mediated transport events (Figures 5B-D), which was in agreement with a recent report in human HEK293T cells (28). Nsp6, Nsp8 and M could also interact with proteins involved in cellular metabolic processes, such as ATP biosynthesis and cellular biosynthetic process (Figures 5B-D), indicating that SARS-CoV-2 might impair the energy supply of human CMs. Canonical signaling pathway enrichment analyses found that Nsp6, Nsp8 and M could all interact with protein factors associated with cell injury signaling, such as coronavirus pathogenesis pathway, viral exit from host cells, calcium and cardiac hypertrophy signaling (Figures 5E-G), suggesting that SARS-CoV-2 infection could induce comprehensive cellular injuries and cardiomyopathy in human CMs. Importantly, by zooming in the protein-protein interaction map and GO enrichment analyses, we found Nsp6, Nsp8 and M interactors were associated with cardiac hypertrophy and mitochondrial dysfunction (Figure 5H), and Nsp8 (Figure 5I) and M (Figure 5J) interactors were functionally related to cardiac arrhythmia. Importantly, Nsp6, Nsp8 or M all interacted with ATPase subunits ATP5A1 and ATP5B in hESC-CMs (Figure 5K), which was further validated by Co-IP Western-blotting assay in the Figure 5L. All these data strongly suggest that Nsp6, Nsp8 and M could likely impair ATP biosynthesis in hPSC-CMs. To confirm it, we quantified the cellular ATP levels in hESC-CMs and hiPSC-CMs. As shown in the Figures 5M-N, Nsp6^OE^, Nsp8^OE^ and M^OE^ in hESC-CMs and hiPSC-CMs could prominently reduce the ATP levels when compared to control CMs. All these data demonstrate the detrimental role of Nsp6, Nsp8 and M interactomes in hPSC-CMs, which lead to impaired ATP homeostasis of hPSC-CMs.

**Figure 5.**
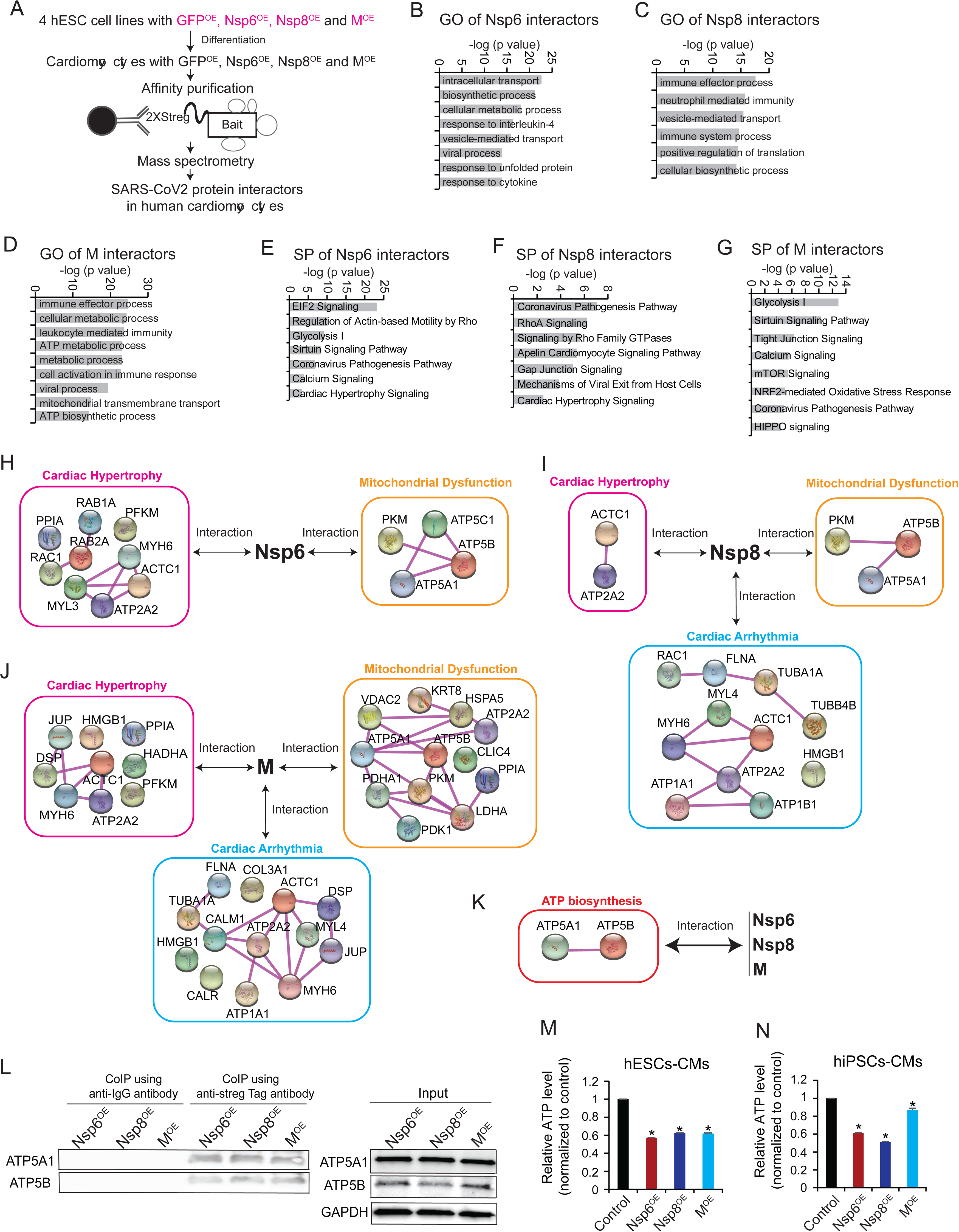
SARS-CoV-2 genes interact with host proteins in hPSC-CMs and reduce cellular ATP level. **(A)** The scheme of Co-IP MS method to globally study SARS-CoV-2 protein interactors in human cardiomyocytes. **(B-D)** Gene Ontology (GO) analyses of Nsp6 interactors (**B**), Nsp8 interactors (**C**) and M interactors (**D**). **(E-G)** Signaling pathways (SP) analyses of Nsp6 interactors (**E**), Nsp8 interactors (**F**) and M interactors (**G**). **(H)** Protein-protein interaction (PPI) shows Nsp6 interactors are associated with cardiac hypertrophy and mitochondrial dysfunction. **(I)** Protein-protein interaction (PPI) shows Nsp8 interactors are associated with cardiac hypertrophy, mitochondrial dysfunction and cardiac arrhythmia. **(J)** Protein-protein interaction (PPI) show M interactors are associated with cardiac hypertrophy, mitochondrial dysfunction and cardiac arrhythmia. **(K)** Protein-protein interaction (PPI) reveals the ATPase subunits ATP5A1 and ATP5B are shared interactors by Nsp6/Nsp8/M proteins in hPSC-CMs. **(L)** Co-IP Western-blotting verification of the interactions of ATP5A1/ATP5B and Nsp6/Nsp8/M proteins in hESCs-CMs. **(M)** ATP level detection in hESCs-derived cardiomyocytes. All bars are shown as mean ± SD. (n=3). A two-tailed unpaired *t*-test was used to calculate *P*-values: \**P* < 0.05 (vs. Control). **(N)** ATP level detection in hiPSCs-derived cardiomyocytes. All bars are shown as mean ± SD. (n=3). A two-tailed unpaired *t*-test was used to calculate *P*-values: \**P* < 0.05 (vs. Control).

### FDA-approved drugs ivermectin and meclizine ameliorate apoptosis of the Nsp6^OE^, Nsp8^OE^ or M^OE^ hPSC-CMs

Nsp6^OE^, Nsp8^OE^ or M^OE^ reduced ATP level of hPSC-CMs, and decreased ATP level has been found to cause apoptosis in various cell types including CMs (29–32). Therefore, we posited that pharmaceutical chemical, which functions to enhance cellular ATP biosynthesis, could potentially reduce apoptosis of Nsp6^OE^, Nsp8^OE^ or M^OE^ hESC-CMs. Two U.S. Food and Drug Administration (FDA)-approved drugs, ivermectin (antiparasitic) and meclizine (antiemetic), were previously reported to protect mitochondrial function and maintain cellular ATP level (33–35). Interestingly, after adding ivermectin (0.5 µM) or meclizine (0.5 µM) into culture media for 3 hrs, the ATP levels in Nsp6^OE^, Nsp8^OE^ and M^OE^ hESC-CMs significantly increased compared to the control groups without treatment (Figure 6A). Notably, after 48 hrs of treatment with ivermectin (0.5 µM) or meclizine (0.5 µM), the ratios of TUNEL^+^ CMs in Nsp6^OE^, Nsp8^OE^ and M^OE^ hiPSC-EBs were significantly decreased compared with control hiPSC-EBs without drug treatment (Figures 6B-F). All these results demonstrate that reduced cellular ATP level is, at least in part, the cause of apoptosis induced by SARS-CoV-2 genes, and FDA-approved drugs ivermectin and meclizine could ameliorate apoptosis of hPSC-CMs via restoring the cellular ATP level.

**Figure 6.**
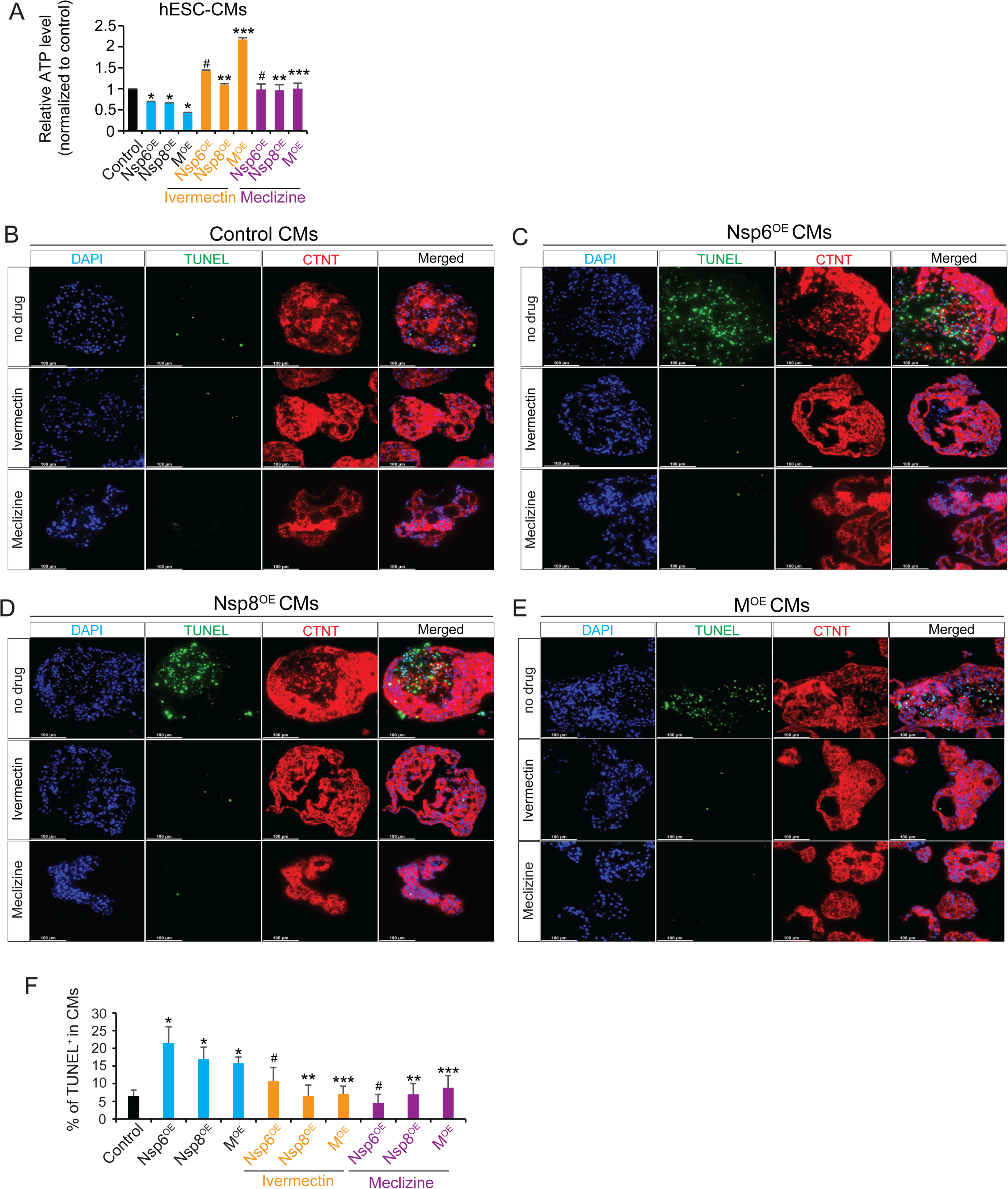
FDA-approved drugs can ameliorate SARS-CoV-2 genes-induced cell death through restoring cellular ATP level. **(A)** ATP level detection in hESC-derived CMs. Ivermectin final concentration is 0.5 µM. Meclizine final concentration is 0.5 µM. Treatment time is 3 hours. **(B-E)** Representative immunostaining images of TUNEL^+^ and CTNT^+^ CMs in control hiPSC-derived EBs **(B)**, hiPSCs-derived EBs with Nsp6 overexpression (Nsp6^OE^) **(C)**, hiPSC-derived EBs with Nsp8 overexpression (Nsp8^OE^) **(D)** and hiPSC-derived EBs with M overexpression (M^OE^) **(E)**. Ivermectin final concentration is 0.5 µM. Meclizine final concentration is 0.5 µM. Treatment time is 2 days. **(F)** Statistical data analysis of immunostaining results from (**B-E**). All bars are shown as mean ± SD. (n=3). A two-tailed unpaired *t*-test was used to calculate *P*-values. \**P* < 0.05 (vs. Control), ^#^*P* < 0.05 (vs. Nsp6^OE^), \*\**P* < 0.05 (vs. Nsp8^OE^), \*\*\**P* < 0.05 (vs. M^OE^).

### Ivermectin and meclizine mitigate SARS-CoV-2 genes-induced abnormal calcium handling and electrical dysfunctions of hPSC-CMs

ATP Homeostasis plays an essential role in heart function. Insufficient ATP level could impair intracellular Ca^2+^ signaling and excitation-contraction coupling of CMs to compromise contractile capacity(36–38). We measured [Ca^2+^]_i_ transients in the spontaneously beating Nsp6^OE^, Nsp8^OE^ or M^OE^ hESC-CMs by using a microfluorimetry. Representative time plots of changes in [Ca^2+^]_i_ were shown in the Figures 7A-B. While control hESC-CMs exhibited regular [Ca^2+^]_i_ transients of uniform amplitudes, [Ca^2+^]_i_ transients in the Nsp6^OE^, Nsp8^OE^ or M^OE^ CMs occurred at slower and irregular rates and developed the delayed aftertransients, which indicated the increased Ca^2+^ leak from sarcoplasmic reticulum (SR). However, adding ivermectin (0.5 µM) or meclizine (0.5 µM) for 1 hr. prominently restored the abnormal rate and rhythmicity of spontaneous [Ca^2+^]_i_ transients and abrogated aftertransients in the Nsp6^OE^, Nsp8^OE^ or M^OE^ hESC-CMs. Additionally, histograms analysis of the interspike intervals (ISIs) generated from [Ca^2+^]_i_ transient recordings revealed a larger range of variations in the ISI distribution from the Nsp6^OE^, Nsp8^OE^ or M^OE^ hESC-CMs than control hESC-CMs, which could be markedly reduced following treatment with ivermectin or meclizine (Figures 7C-D).

**Figure 7.**
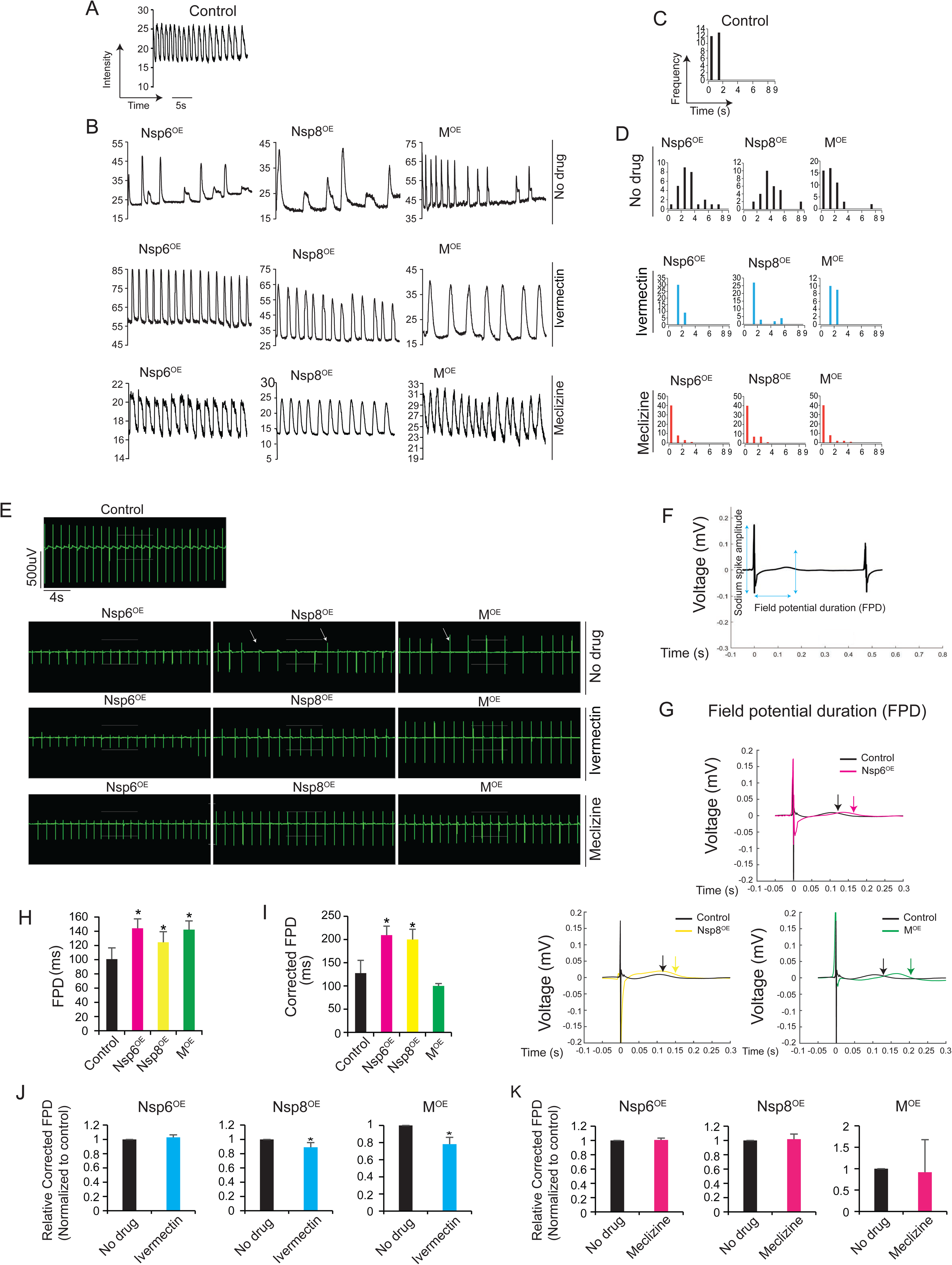
SARS-CoV-2 genes-induced abnormal calcium handling and electrical dysfunction in human cardiomyocytes can be ameliorated by FDA-approved drugs. **(A)** Calcium handling analysis in control hESC-CMs. Y-axis represents intensity and X-axis represents time (in second, s). **(B)** Calcium handling analysis in Nsp6^OE^, Nsp8^OE^ or M^OE^ hESC-CMs treated without or with drugs for 1 hour. Y-axis means intensity. X-axis represents time (in second, s). **(C)** Interspike interval (ISI) distribution analysis from (**A**). Y-axis means beating frequency in a specific time window. X-axis means time (in second, s). **(D)** Interspike interval (ISI) distribution analysis from (**B**) in different CM cell lines treated with or without drugs for 1 hour. **(E)** Electrophysiology analysis by multi-electrode arrays (MEAs) in hESC-CMs. Representative field potential recordings of spontaneous beating hESC-CMs. For drug treatment assay, the final concentrations of ivermectin and meclizine were 0.5 µM. The data was acquired after 1 hour of drug treatment. **(F)** An illustration of the field potential duration (FPD, ms) from hESC-CMs. **(G)** Representative traces illustrate the field potential duration (FPD, ms) from control and Nsp6^OE^, Nsp8^OE^ and M^OE^ hESC-CMs, respectively (from left to right). **(H)** Field potential duration (FPD) analysis of MEAs data in hESC-CMs. All bars are shown as mean ± SD. (n = 5). A two-tailed unpaired t-test was used to calculate *P*-values. \**P* < 0.05 (vs. Control). **(I)** Corrected FPD analysis of MEA data in hESC-CMs. Quantification of field potential duration (FPD) was corrected by beat rate (FPDc) in spontaneous experiments using the Bazett formula. All bars are shown as mean ± SD. (n= 5). A two-tailed unpaired t-test was used to calculate P-values. \**P* < 0.05 (vs. Control). **(J)** Corrected FPD (FPDc) analyses of MEA data in hESC-CMs treated with ivermectin for 1 hour. All ivermectin-induced FPDc were normalized to that of control without ivermectin treatment. All bars are shown as mean ± SD, (n=3). A two-tailed unpaired t-test was used to calculate P-values. \**P* < 0.05 (ivermectin vs. no drug). **(K)** Corrected FPD (FPDc) analyses of MEA data in hESC-CMs treated with meclizine for 1 hour. All meclizine-induced FPDc were normalized to that of control without meclizine treatment. All bars are shown as mean ± SD, (n=3). A two-tailed unpaired t-test was used to calculate P-values. \**P* < 0.05 (Meclizine vs. no drug).

Next, we performed the multiple-electrodes arrays (MEAs) to capture the filed potential and contractions of hESC-CMs. As shown in the Figure 7E, field potential recording from control hESC-CMs revealed regular contractions, whereas Nsp6^OE^, Nsp8^OE^ or M^OE^ hESC- CMs exhibited irregular and slower beating patterns. Administration of ivermectin or meclizine effectively restored beating rate and rhythmicity of Nsp6^OE^, Nsp8^OE^ and M^OE^ hESC-CMs back to a similar level as control CMs (Figure 7E). We then measured the field potential duration (FPD) of beating hESC-CMs to assess CM repolarization (Figure 7F). Nsp6^OE^, Nsp8^OE^ and M^OE^ hESC-CMs exhibited elongated FPDs than control hESC-CMs (Figure 7G, three panels; Figurer 7H). After adjustment of FPD for beating rate using the Bazett’s formula, the corrected FPD (FPD_c_) of CMs with Nsp6^OE^and Nsp8^OE^, but not M^OE^, remained significantly prolonged compared to control CMs (Figure 7I), suggesting that Nsp6 and Nsp8 altered repolarization properties of hESC-CMs. These MEAs results, together with abnormal calcium handling, indicate that SARS-CoV-2 gene could disrupt the mechanisms underlying pacemaking and repolarization of human CMs, which could possibly contribute to cardiac arrhythmia in COVID-19 patients. Furthermore, ivermectin significantly reduced FPD_c_ in Nsp8^OE^ and M^OE^ CMs (Figure 7J), whereas no significant effect of meclizine on FPD_c_ was observed (Figure 7K).

## Discussion

Understanding the cardiac manifestations in patients with SARS-CoV-2 infection is critical for health care of acute and post-acute COVID-19 patients (39, 40). In this study, we reveal the genome-wide responses of host hPSC-CMs to SARS-CoV-2 viral genes Nsp6, Nsp8 and M in whole transcriptome and interactome. Our findings uncover the compromised cellular ATP level of hPSC-CMs which overexpress Nsp6, Nsp8 or M. FDA-approved drugs, ivermectin and meclizine, could sustain the cellular ATP level to attenuate apoptosis and dysfunctions of hPSC-CMs upon Nsp6, Nsp8 or M overexpression. Our results suggest that the impaired ATP hemostasis might play a vital role in SARS-CoV-2 gene-induced CM injury in the heart, and possibly in other organs/tissues that are targets of SARS-CoV-2 as well. Notably, although both Nsp6^OE^, Nsp8^OE^ and M^OE^ and SARS-CoV-2 infection led to concordant transcriptomic changes of hPSC-CMs (Figures 3B-C), ∼70% DEGs were solely induced by SARS-CoV-2 virus (Figure 3G-H), suggesting that the other SARS-CoV-2 genes might also affect the transcriptome of host human CMs via different targets or under different mechanisms. Since the unique pathways induced by SARS-CoV-2 virus were related to apoptosis, gene transcription and multiple metabolic processes, these results imply that at least some of other SARS-CoV-2 genes might also contribute to CM injury and affect the metabolism of host human CMs.

Multiple clinical studies reported that myocardial dysfunctions and injuries were commonly found in COVID-19 patients, and positively contributed to the overall mortality (3-5,22,41). Recently, Bailey et al.(9) reported that SARS-CoV-2 can directly and specifically infect CMs within the hearts of COVID-19 patients. Myocardial tissues from 4 autopsy of COVID-19 patients with clinical myocarditis were examined. In all autopsies, CM infection was identified by intramyocyte expressions of viral RNA and protein, and macrophage infiltration associated with areas of myocyte cell death. This study provided direct evidence for CM infection in the COVID-19 patient hearts. Additionally, several research groups reported that SARS-CoV-2 virus could infect human ESC/iPSC-derived CMs, and induce apoptosis (15, 16), sarcomere fragmentations (17) and electrical and mechanical dysfunctions (18) compared with mock control. After infection, the intramyocyte viral particles were observed by using transmission electron microscopy. All these studies indicate that hPSC-CMs could be utilized as an *in vitro* model for conducting mechanistic studies of SARS-CoV-2 viral genes in human CMs. In this study, we identified the susceptibility of hPSC-CMs to individual SARS-CoV-2 genes. Nsp6 and Nsp8 are the non-structural proteins of SARS-CoV-2. M is the structural protein of SARS-CoV-2, which is the most abundant structural protein in the virus particle. The mechanisms of CM death induced by those genes have not been previously described. Interestingly, we found that enforced expression of Nsp6, Nsp8 or M was sufficient to induce apoptosis and dysfunction of hPSC-CMs (Figures 4 and 7A,B,E), which phenocopied SARS-CoV-2 infected hPSC-CMs from other reports (9,15,17,42). Whole mRNA-seq revealed global transcriptional changes of Nsp6^OE^, Nsp8^OE^ and M^OE^ hESC-CMs compared to control hESC-CMs (Figures 2C-E), particularly with differentially expressed genes enriched into activated cellular injury and immune responses whereas reduced calcium/gap junction signaling (Figure 3H). Notably, these global transcriptome changes are consistent with the previous published whole mRNA-seq results from SARS-CoV-2 virus directly infected hPSC-CMs (9). These results thus indicate that expression of the exogenous SARS-CoV-2 viral genes could profoundly disrupt the gene expression programs of human CMs, which might subsequently cause CM abnormalities in COVID-19 patients.

The interactions of SARS-CoV-2 viral genes with host cell proteins also play a critical role in CM abnormalities. In this study, by studying the interactome of Nsp6, Nsp8 or M within hESC-CMs, we found that they all interacted with ATPase subunits and compromised the cellular ATP level in hPSC-CMs (Figures 5A,K). Bailey et al. also found downregulated ATP metabolic process in SARS-CoV-2 infected hPSC-CMs compared to control using whole mRNA-seq (9). Therefore, our findings suggest a possible central role of ATP homeostasis in SARS-CoV-2-induced tissue injuries in CMs, and highly likely in other SARS-CoV-2 vulnerable tissue cells in lung or kidney, although the mechanisms how Nsp6, Nsp8 or M could highjack ATPase remain to be elucidated.

Due to the consistent contractions, heart muscle cells consume a large amount of energy, which makes CMs one of the most vulnerable cell types to insufficient ATP supply. Therefore, we explored the pharmaceutical strategies to enhance cellular ATP levels, and found two FDA-approved drugs ivermectin and meclizine significantly reduced SARS-CoV-2 genes-induced cell death and electrical dysfunctions in human CMs. Ivermectin belongs to the group of avermectins (AVM), which is a series of 16-membered macrocyclic lactone compounds discovered in 1967. FDA approved it for human use in 1987. Although ivermectin is used for treating parasitic infection, it was identified as a mitochondrial ATP protector in CMs and increased mitochondrial ATP production in human CMs (33), which was verified in our study as well. Interestingly, a study reported that ivermectin was an inhibitor of SARS-CoV-2 *in vitro*, and a single treatment for 48 hrs. led to ∼5000-fold reduction of virus in cell culture (34). Notably, a recent clinical study reported that administration of ivermectin was associated with significantly lower mortality in hospitalized COVID-19 patients(43), although it is unclear whether ivermectin could reduce mortality via preserving ATP levels in SARS-CoV-2 vulnerable heart muscle and/or other tissue cells. Meclizine, an over-the-counter anti-nausea and -dizziness drug, was identified via a ’nutrient-sensitized’ chemical screen. Meclizine is primarily used as an antihistamine. However, it has been reported that meclizine had cardio-protection effect through promoting glycolysis of CMs (35), which increased ATP synthesis. Other study also reported that meclizine stimulated glycolysis, mitigated ATP depletion and protected mitochondria function (44). Those studies support our findings that meclizine might increase ATP level to mitigate SARS-CoV-2 gene-induced cell death of hPSC-CMs. Currently, whether administration of meclizine is associated with reduced mortality in hospitalized COVID-19 patients is still unknown and requires further clinical investigations.

## Acknowledgements

We thank the Proteomics Core of Indiana University School of Medicine (IUSM) for Co-IP-MS. Work in the IUSM Proteomics Core was supported, in part, with support from the Indiana Clinical and Translational Sciences Institute, which is funded by the UL1TR002529 from the National Institutes of Health, National Center for Advancing Translational Sciences, Clinical and Translational Sciences Award. Acquisition of the IUSM Proteomics core instrumentation used for this project, the Orbitrap Fusion Lumos, was provided by the Indiana University Precision Health Initiative. We also thank Indiana Center for Biological Microscopy of Indiana University School of Medicine (IUSM) for cell imaging.

## Author contributions

JL and LY initiated and designed studies. JL performed all experiments and data analyses. SW, LH, CW, ES, YW and YLL assisted in whole RNA-seq. YZ and JW assisted in bioinformatics. YL, WS and ML supported calcium handling and MEA experiments and data analyses. JL, JW, ML and LY wrote the manuscript.

## Abbreviations and Acronyms

SARS-CoV-2: severe acute respiratory syndrome-coronavirus-2
COVID-19: coronavirus disease-2019
ACE2: angiotensin converting enzyme 2
hPSCs: human pluripotent stem cells

---

**Supplementary Figure 1.**
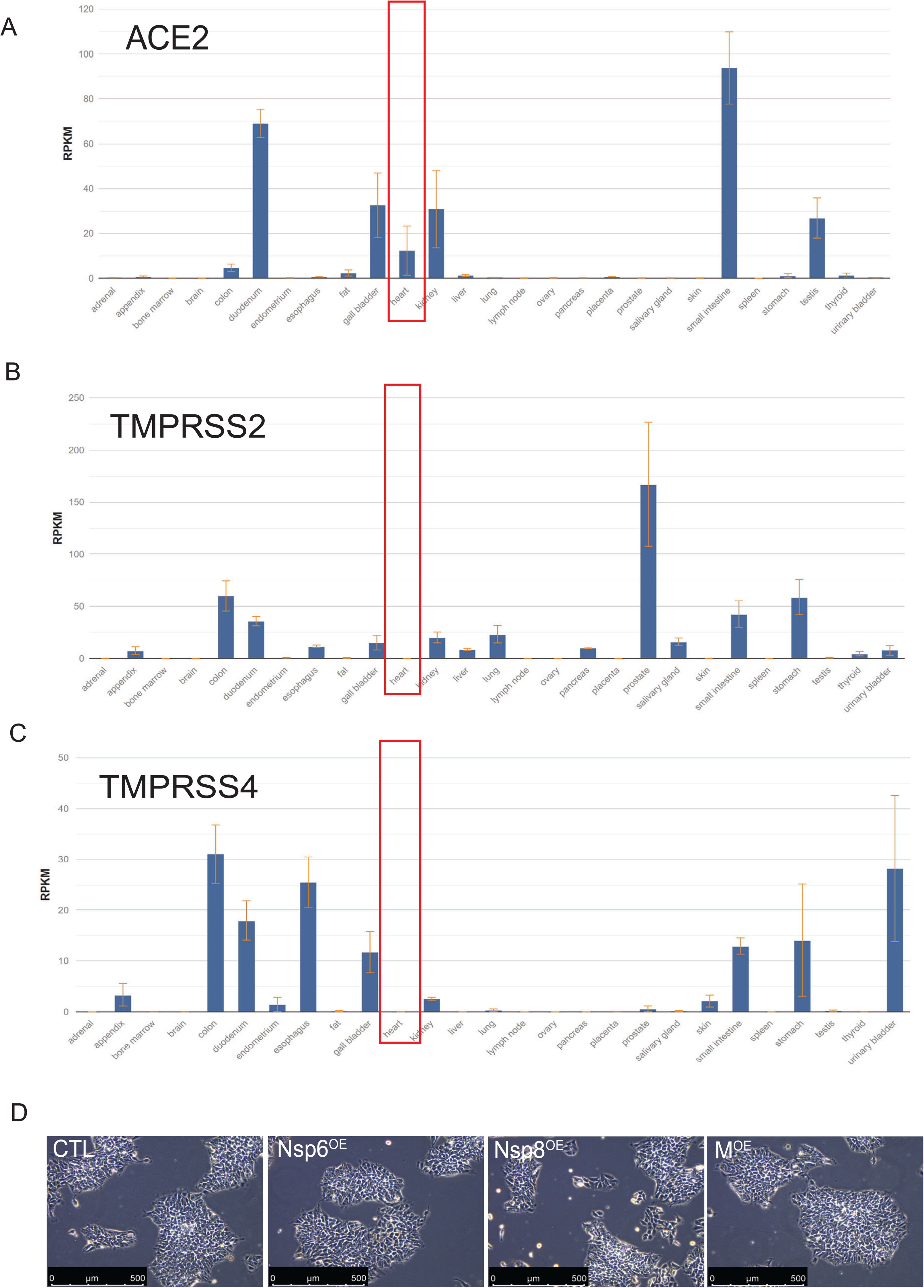
Expression of SARS-CoV-2 receptor genes in human heart. (A) RNA expression level of ACE2 gene in adult human heart tissues from RNA-seq data (NCBI database). (B) RNA expression level of TMPRSS2 gene in adult human heart tissues from RNA-seq data (NCBI database). (C) RNA expression level of TMPRSS4 gene in adult human heart tissues from RNA-seq data (NCBI database). (D) Representative images of control and Nsp6OE, Nsp8OE and MOE in hESC lines.

**Supplementary Figure 2.**
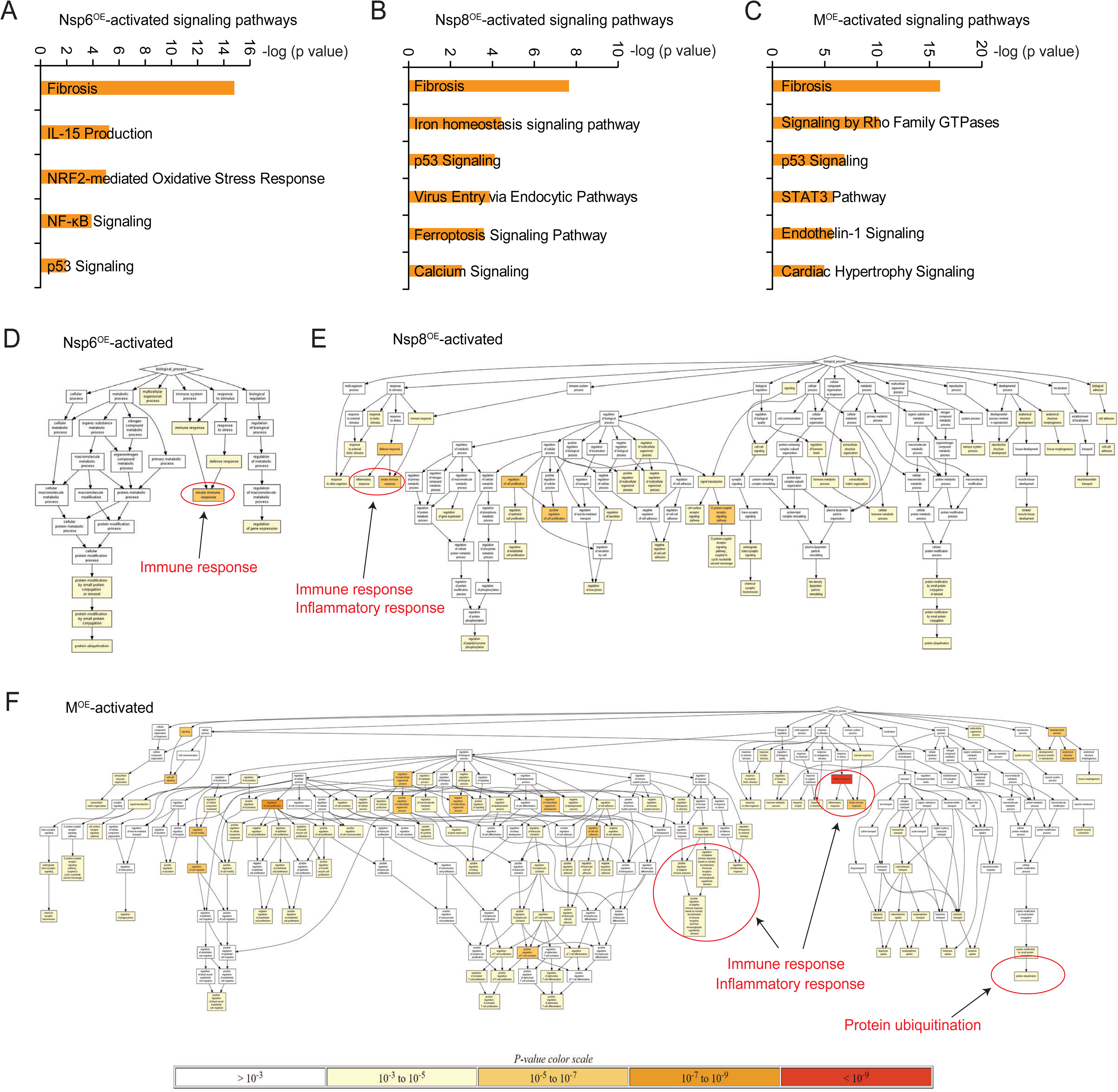
MRNA-seq reveals that SARS-CoV-2 genes induces cell injuries and dysfunction in hESC-CMs. (A-C) Signaling pathways activated by Nsp6OE, Nsp8OE and MOE in hESC-CMs. (D-F) GO Hierarchy activated by Nsp6OE, Nsp8OE and MOE in hESC-CMs.

**Supplementary Figure 3.**
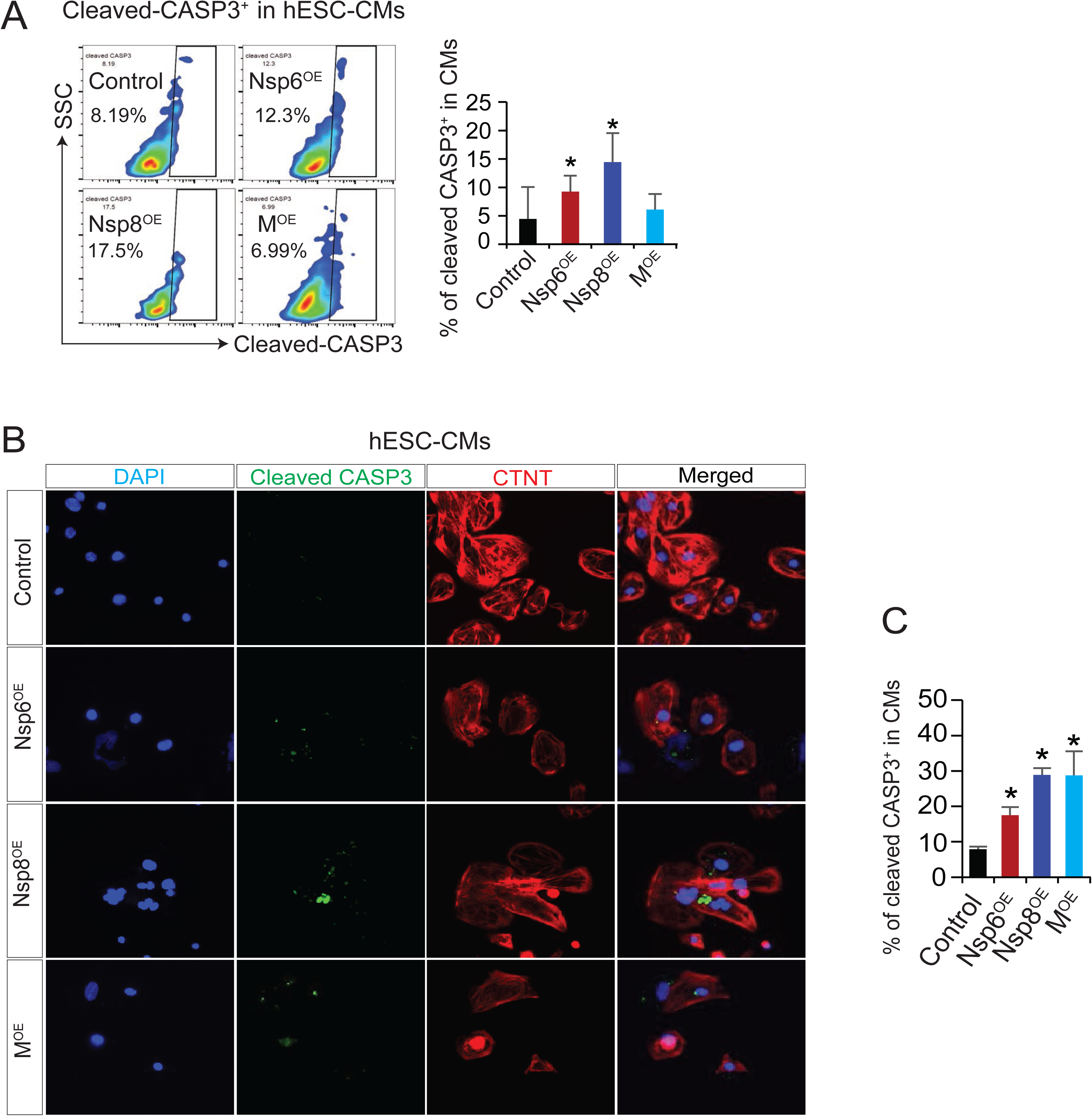
Overexpression of SARS-CoV-2 genes can cause cell death in hPSC-CMs. (A) Flow cytometry analysis of cleaved CASP3+ cells in hESCs-derived cardiomyocytes (CMs) with overexpression of SARS-CoV-2 genes. All bars are shown as mean ± SD. (n=8). A two-tailed unpaired t-test was used to calculate P-values: *P < 0.05 (vs. Control). (B) Immunofluorescence of cleaved CASP3+ and CTNT+ cells in hESC-CMs with overexpression of SARS-CoV-2 genes. (C) Statistical data analysis of immunostaining from (B). All bars are shown as mean ± SD. (n=3). A two-tailed unpaired t-test was used to calculate P-values: *P < 0.05 (vs. Control).

## Supplemental Information

### Materials and Methods

#### Human pluripotent stem cells culture and cardiomyocytes differentiation

Human embryonic stem cell (hESC) line H9 and human induced pluripotent stem cell (hiPSC) line S3 were cultured on Matrigel (BD Biosciences)-coated plates in mTesR1 medium ^1,2^. For cardiomyocyte differentiation, the monolayer differentiation method was used based on published protocol ^3^ with minor modification. Briefly, differentiation was initiated by removing the mTeSR1 medium and adding RPMI1640/B27 (no insulin) medium with 6 μM CHIR99021 from day 0 to day 1. Then cells were induced in only RPMI1640/B27 (no insulin) medium from day 1 to day 3, followed with induction in RPMI1640/B27 (no insulin) medium with 5 μM XAV-939 from day 3 to day 5. After day 5, cells were maintained in RPMI1640/B27 (no insulin) medium without any chemicals. Beating cardiomyocytes could be observed after differentiation of 7 days. For embryoid body (EBs) differentiation (3D differentiation), hiPSCs were differentiated towards CMs using the previously established protocol ^4,5^. Briefly, cardiomyocyte differentiation was conducted with EBs formation. EBs were treated with StemPro-34 SFM (1X) medium (10639011, Gibco.) with the following conditions: days 0-1 with BMP4 (2.5 ng/ml); days 1-4 with BMP4 (10 ng/ml), FGF-2 (5 ng/ml) and Activin A (2 ng/ml); and days 4-20 with XAV939 (5µM), after day 4 the medium was changed every 4 days. Beating EBs could be observed at day 10-13 after differentiation. For drug treatment assay, all beating EBs were maintained in DMEM medium (no glucose, Gibco) with 10% FBS. Both of the final concentration of Ivermectin and Meclizine was 0.5µM (drugs were dissolved in DMSO). The treatment time for apoptosis assay was 2 days. All cytokines were from R&D Systems. All chemicals or drugs were from Sigma Aldrich.

#### Lentivirus production and cell transduction

The lentiviral vector pLVX-EF1alpha-IRES-Puro-2xStrep-SARS-CoV-2-GFP (Nsp6, Nsp8, M or Orf9c) was transfected into the HEK293T cells with packaging plasmids psPAX2 and pMD2.G (gifts from Dr. Guang Hu in NIH) using X-treme GENE 9 transfection reagent (Roche) according the transfection manual. After transfection of 2 days, viral supernatant was collected and cellular debris was removed by syringe filtering (0.45 μm pore size; Millipore). H9 hESCs or S3 hiPSCs cultured in mTesR1 medium were incubated with virus media for 4h, followed with fresh mTesR1 medium culture for overnight. Same infection was repeated after 24 hr. Puromycin was added to select puromycin-resistant H9 hESCs or S3 hiPSCs clones after 48h virus infection.

#### RT-qPCR

RNAs were isolated by RNeasy Kit (Qiagen). CDNA synthesis was carried out using High-Capacity RNA-to-cDNA™ Kit (Applied Biosystems). RT-qPCR was performed on QuantStudio 6 and 7 Flex Real-Time PCR Systems with Fast SYBR Green Master Mix (Applied Biosystems) according to the manufacturer’s instructions. The RT-qPCR analysis were normalized to internal control GAPDH or beta-ACTIN using 2^−ΔΔCt^ method^6^. RT-qPCR data were presented with mean ± S.D. from at least three independent experiments. For all primers’ information, please see the Table S3 Oligonucleotides.

#### Western Blotting

Proteins were extracted by Complete™ Lysis-M EDTA-free kit (Roche, 04719964001). Protein samples were mixed with NuPAGE™ LDS Sample Buffer (4X), boiled for 5 min at 95 °C, run on 4–15% Mini-PROTEAN TGX Gels (Bio-Rad), transferred to PVDF membrane by using Trans-Blot® Turbo™ Transfer System (Bio-Rad), blocked with 1X TBST buffer containing 10% not-fat milk, and incubated overnight at 4 °C with the corresponding primary antibodies with 5% BSA in 1X TBST buffer. Membranes were washed three times for 5 min each with 1X TBXT buffer, and then incubated for 1 h at room temperature with horseradish-peroxidase-conjugated secondary antibodies in blocking solution with 5% BSA in 1X TBST buffer. Membranes were washed again three times for 5 min with 1X TBST before revealing them with ECL Western Blotting Substrate (Pierce) and ChemiDoc Imaging System (Bio-Rad).

#### Co-immunoprecipitation (Co-IP)

Cell proteins were extracted by Pierce™ Classic Magnetic IP/Co-IP Kit (Thermo Scientific, 88804) according to the manuals. Protein Co-immunoprecipitation (Co-IP) was performed by using Pierce™ Classic Magnetic IP/Co-IP Kit as well. Co-IP protein samples analysis were performed by using Western blotting or submitted to Proteomics Core Facility at the Indiana University School of Medicine (IUSM) for mass spectrometry analysis.

#### Flow cytometry

Flow cytometry was performed based on our previous protocol ^7^. Briefly, attached cells were harvested and dissociated by using 0.25% trypsin-EDTA at 37°C incubator for 10 min. The dissociated cells were fixed in 4% PFA (diluted with 16% Paraformaldehyde (formaldehyde) aqueous solution) at room temperature for 10 min and washed 3 times with 1X PBS. Cells were incubated in 1X blocking PBS buffer (containing 2% goat serum or 5% BSA plus 0.1% saponin) with corresponding primary antibodies at 37°C for 1h, following with corresponding secondary antibodies staining at 37°C for 1 h. For TUNEL (Terminal deoxynucleotidyl transferase dUTP nick end labeling) experiments, staining was carried out by using In Situ Cell Death Detection Kit, Fluorescein (11684795910 Roche) according to the manual. Flow cytometry analysis was performed on Attune NxT Flow Cytometer (Thermo Fisher Scientific). Data were analyzed by using FlowJo software (Treestar).

#### SARS-CoV-2 spike pseudovirus infection

The SARS-CoV-2 Spike Pseudotyped Lentivirus with GFP reporter (60 concentration, 60X) was purchased from Virongy (USA). HESCs were cultured in mTesR1 medium on 6-well plate coated with Matrigel. Cells were infected with 1X or 2X SARS-CoV-2 Spike Pseudotyped Lentivirus. After 48 hours of infection, flow cytometry was performed to detect GFP^+^ cells. Cells without virus infection were the blank control cells. Flow cytometry analysis was performed on Attune NxT Flow Cytometer (Thermo Fisher Scientific, USA). Data were analyzed by using FlowJo software (Treestar, USA).

#### Apoptosis assay of live cells

Live cells apoptosis analysis was performed using Annexin-V-FLUOS Staining Kit (11858 777001, Roche) according to the manual. Briefly, cells were harvested and dissociated by using 0.25% trypsin-EDTA at 37°C incubator for 10 min. Single cells were washed in 1X PBS, then resuspend and incubated in 100 µl of Annexin-V-FLUOS labeling solution for 15 min at room temperature, followed with the analysis on BD LSRII cytometer (Becton Dickinson) or Attune NxT Flow Cytometer (Thermo Fisher Scientific). Data were analyzed by using FlowJo software (Treestar).

#### ATP detection assay

The level of ATP within the cells was measured by Luminescent ATP Detection Assay Kit (ab113849, Abcam) according to the manual. Briefly, flow cytometry or hemocytometer was used to adjust live cell number. Same cell number for each group was used for the ATP detection. Live cells were resuspended in 50μl detergent solution with 1200 rpm shaking on Eppendorf ThermoMixer C for 5 min at room temperature. Then 50μl substrate solution was added to the detergent solution with 1200 rpm shaking on Eppendorf ThermoMixer C for 5 min at room temperature. All solution was transferred to Nunc™ MicroWell™ 96-Well, (Nunclon Delta-Treated, Flat-Bottom Microplate, White Polystyrene Plate, Thermo Scientific, 136101), followed with luminescence detection on GloMax Discover Microplate Reader (GM3000, Promega). For drug treatment assay, both of the final concentration of Ivermectin and Meclizine was 0.5µM, the treatment time was 3h.

#### Intracellular calcium handling

Cardiomyocytes were attached on 12-well plate coated with Matrigel and loaded with X-Rhod-1 (X14210, Invitrogen) in DMEM medium (no glucose) with 10% FBS containing Pluronic F-127 (final concentration 0.02%, P2443, Sigma) for 15 min at 37 °C incubator. Videos were acquired at a rate of 50 frames per second using the All-in-One Fluorescence Microscope BZ-X800 (KEYENCE CORPORATION). For drug treatment assay, both of the final concentration of Ivermectin and Meclizine were 0.5 µM (drugs were dissolved in DMSO). The calcium handling signaling was acquired after 1 hour of drug treatment. Videos were analyzed in ImageJ software using a custom script that calculated the temporal changes in calcium fluorescence intensity.

#### Multi-electrode arrays (MEAs) for cardiomyocytes

Electrophysiology experiments were performed for cardiomyocytes after day 30 differentiation. All experiments were performed in DMEM medium (no glucose, Gibco) with 10% FBS. Cardiomyocytes were attached on cytoview MEA 24-well white plate (M384-tMEA-24W-5, Axion Biosystems) coated with Matrigel. Electrophysiology data were acquired by the Maestro Edge MEA System (Axion Biosystems) with the integrated environment chamber controlling the heat at 37 degree and the CO₂ at 5%.

#### Immunofluorescence

For immunostaining of attached cells, cells were fixed with 4% PFA (diluted from 16% Paraformaldehyde aqueous solution) for 10 min at room temperature. After washing with 1X PBS, cells were blocked for 1 h with 1X PBS blocking buffer containing 2% goat serum (or 5% BSA) and 0.1% saponin. Staining with corresponding primary antibodies diluted with blocking buffer was performed at -4°C for overnight. Staining with secondary antibodies were performed on next day, following with nucleus staining with DAPI. For beating EBs immunostaining, live EBs were fixed with 4% PFA (diluted from 16% Paraformaldehyde aqueous solution) for 10 min at room temperature, followed with 15% sucrose solution at room temperature until all EBs sunk. Then EBs were embed in OCT and cryosectioning was performed. The EBs immunostaining protocol was the same as that of attached cells. For TUNEL (Terminal deoxynucleotidyl transferase dUTP nick end labeling) experiments, staining of attached cells or EBs sections were carried out using In Situ Cell Death Detection Kit, Fluorescein (11684795910 Roche) according to the manual. Leica DM6B image system was used for imaging.

#### Co-immunoprecipitation mass spectrometry (CoIP-MS)

Beads were submitted to the proteomics core where they were covered in 8 M Urea, 50 mM Tris-HCl, pH 8.5, reduced with 5 mM tris (2-carboxyethyl) phosphine hydrochloride (TCEP) at room temperature for 30 min and alkylated with 10 mM chloroacetamide (CAM) for 30 min in the dark at room temperature. Digestion was carried out using Trypsin/Lys-C Mass spec grade protease mix (Promega, V5072) at a 1:100 protease to substrate ratio, overnight at 37 °C. The reaction was quenched with 0.5 % formic acid prior to LC-MS. Samples were analyzed using a 5 cm trap column and 15 cm (2 µm particle size, 50 µm diameter) EasySpray (801A) column on an UltiMate 3000 HPLC and Q-Exactive Plus mass spectrometer (Thermo Fisher Scientific). Solvent B was increased from 5%-28% over 155 min, to 35% over 5 min, to 65% over 10 min and back to 5% over 1 2 min(Solvent A: 95% water, 5% acetonitrile, 0.1% formic acid; Solvent B: 100% acetonitrile, 0.1% formic acid). A data dependent top 20 method acquisition method was used with MS scan range of 350-1600 m/z, resolution of 70,000, AGC target 3e6, maximum IT of 50 ms. MS2 settings of fixed first mass 100 m/z, normalized collision energy of 36, isolation window of 1.5 m/z, resolution of 35,000, target AGC of 1e5, and maximum IT of 250 ms. For dd acquisition a minimum AGC of 2e3 and charge exclusion of 1, and ≥7 were used.

#### Tandem mass tag-mass spectrometry (TMT-MS)

##### Cell and tissue preparation

Cells were lysed in 8 M urea, 50 mM Tris-HCl, pH 8.5. Samples were sonicated in a Bioruptor® sonication system from Diagende Inc. (30 sec/30 sec on/off cycles for 15 minutes, 4°C). Following centrifugation at 12,000 rpm for 15 minutes, protein concentrations were determined using a Bradford protein assay (cat. num. 5000002, Bio-Rad). Protein samples in equal amounts (30 μg) were reduced with 5 mM TCEP and alkylated with 10 mM (CAM). Samples were diluted with 100 mM Tris-HCl to a final urea concentration of 2 M and digested overnight with Trypsin/Lys-C Mix Mass Spectrometry (1:100 protease : substrate ratio, cat. num. V5072, Promega).

##### Peptide purification and labeling

Peptides were desalted on 50 mg Sep-Pak® Vac (Waters Corporation) employing a vacuum manifold. After elution from the column in 70% acetonitrile (ACN) 0.1% formic acid (FA), peptides were dried by speed vacuum and resuspended in 24 µL of 50 mM triethylammonium bicarbonate (TEAB). Peptide concentration was measured using Pierce Quantitative Colorimetric Peptide Assay Kit (cat. num. 23275, Thermo Fisher Scientific) to ensure that an equal amount of each sample was labeled. Samples were then Tandem Mass Tag (TMT) labeled with 0.2 mg of reagent resuspended in 20 µL acetonitrile for two hours at room temperature (WT:127N, Nsp6:128N, Nsp8:129N, M:130N, Orf9c:131; cat. num. 90309, Thermo Fisher Scientific TMT10plex™ Isobaric Label Reagent Set; lot no. UH285567 and 131C lot UD280157A). Labelling reactions were quenched with hydroxylamine at room temperature 15 minutes. Labelled peptides were then mixed and dried by speed vacuum.

##### High pH basic fractionation

The peptide mixture was resuspended in 0.1% TFA (trifluoroacetic acid) and fractionated on a PierceTM High pH reversed-phase peptide fractionation spin column using vendor methodology (cat. num. 84868). Each fraction was dried by speed vacuum and resuspended in 30 µL 0.1% FA.

##### Nano-LC-MS/MS Analysis

Nano-LC-MS/MS analyses were performed on an EASY-nLC™ HPLC system coupled to an Orbitrap Fusion™ Lumos™ mass spectrometer (Thermo Fisher Scientific). Half of each fraction was loaded onto a reversed phase PepMap™ RSLC C18 column with Easy-Spray tip at 400 nL/min (ES802A, 2 μm, 100 Å, 75 μm x 25 cm). Peptides were eluted from 4-33% B over 120 minutes, 33%-80% B over 5 mins, and dropping from 50-10%B over the final 4 min min (Mobile phases A: 0.1% FA, water; B: 0.1% FA, 80% Acetonitrile). Mass spectrometer settings include capillary temperature of 300 °C and ion spray voltage was kept at 1.9 kV. The mass spectrometer method was operated in positive ion mode with a 4 second cycle time data-dependent acquisition with advanced peak determination and Easy-IC on (internal calibrant). Precursor scans (m/z 375-1600) were done with an orbitrap resolution of 120000, 30% RF lens, 105 ms maximum inject time (IT), standard automatic gain control (AGC) target. MS2 filters included an intensity threshold of 2.5e-4, charges states of 2 to 6, 70% precursor fit threshold, and 60 s dynamic exclusion with dependent scan being performed on only one charge state per precursor. Higher-energy collisional dissociation (HCD) MS2 scans were performed at 50k orbitrap resolution, fixed collision energy of 37%, 200% normalized AGC target, and dynamic maximum IT.

##### Data analysis

Resulting RAW files were analyzed in Proteome Discover 2.4 (Thermo Fisher Scientific) with FASTA databases including Swiss-Prot UniProt Homo sapiens sequences (downloaded 09/17/2019) plus common contaminants. SEQUEST HT searches were conducted with a maximum number of 2 missed cleavages; precursor mass tolerance of 10 ppm; and a fragment mass tolerance of 0.02 Da. Static modifications used for the search were, 1) carbamidomethylation on cysteine (C) residues; 2) TMT sixplex label on lysine (K) residues and the N-termini of peptides (for TMT quant samples only). Dynamic modifications used for the search were oxidation of M, phosphorylation on S, T, Y, and acetylation of N-termini. IP-MS Sequest results were imported into Scaffold (Proteome Software) for Fishers exact test comparison. TMT quantification methods utilized isotopic impurity levels available from Thermo Fisher. Percolator False Discovery Rate was set to a strict setting of 0.01 and a relaxed setting of 0.05. Values from both unique and razor peptides were used for quantification. In the consensus workflow, peptides were normalized by total peptide amount with no scaling. Resulting abundance values for each sample, and abundance ratio values from Proteome Discoverer™ were exported to Microsoft Excel and are available in a supplemental file.

#### mRNA-seq

##### KAPA mRNA HyperPrep methods for mRNA sequencing

Total RNA will be first evaluated for its quantity, and quality, using Agilent Bioanalyzer 2100. For RNA quality, a RIN number of 7 or higher is desired. One hundred nanograms of total RNA will be used. cDNA library preparation includes mRNA purification/enrichment, RNA fragmentation, cDNA synthesis, ligation of index adaptors, and amplification, following the KAPA mRNA Hyper Prep Kit Technical Data Sheet, KR1352 – v4.17 (Roche Corporate). Each resulting indexed library will be quantified and its quality accessed by Qubit and Agilent Bioanalyzer, and multiple libraries pooled in equal molarity. The pooled libraries will then be denatured, and neutralized, before loading to NovaSeq 6000 sequencer at 300pM final concentration for 100b paired-end sequencing (Illumina, Inc.). Approximately 30-40M reads per library is generated. A Phred quality score (Q score) is used to measure the quality of sequencing. More than 90% of the sequencing reads reached Q30 (99.9% base call accuracy).

##### Data analysis

Quality control for raw mRNA-seq data was generated by FastQC v0.11.5 (http://www.bioinformatics.babraham.ac.uk/projects/fastqc/). Illumina adapter sequences and low-quality bases were trimmed by Trim Galore v0.4.5 (http://www.bioinformatics.babraham.ac.uk/projects/trim_galore/), followed by sequence mapping of high-quality paired-end reads to human genome (hg38) with the aligner STAR v2.7.2b^8^. We further used bam-filter in ngsutilsj v0.4.8 (https://compgen.io/ngsutilsj) to keep only properly and uniquely mapped paired reads (MAPQ ≥ 10) for downstream analysis. FeatureCounts from package subread v1.6.5^9^ was employed to summarize gene expression levels based on mapped reads according to GENECODE v31 annotation. Analysis of differential expression genes (DEGs) was performed by edgeR v3.32.1^10^, with read counts normalized by the trimmed mean of M-values (TMM) method after lowly expressed genes filtered out by the filterByExpr function using default settings. DEGs due to overexpression of SARS-CoV-2 viral coding genes were identified if their FDR-adjusted *p*-values were less than 0.05 based on the comparison between the viral gene overexpression and the control.

##### Functional enrichment analysis

The functional enrichment analysis of gene ontology (GO) biological process was carried out by using DAVID (https://david.ncifcrf.gov/), THE GENE ONTOLOGY RESOURCE (http://geneontology.org/) and Gorilla (http://cbl-gorilla.cs.technion.ac.il/). Canonical signaling pathway and toxicity analysis were performed on QIAGEN Ingenuity Pathway Analysis (IPA).

##### Protein-protein interaction (PPI) analysis

Protein-Protein Interaction Networks analysis were carried out on STRING software (https://string-db.org/).

##### Quantification and Statistical Analysis

Data comparisons between two groups (gene overexpression versus control) were conducted using an unpaired two-tailed *t*-test. All data were presented as mean ± S.D. from at least three independent experiments. Differences with *P* values less than 0.05 were considered significant.

## KEY RESOURCES TABLE

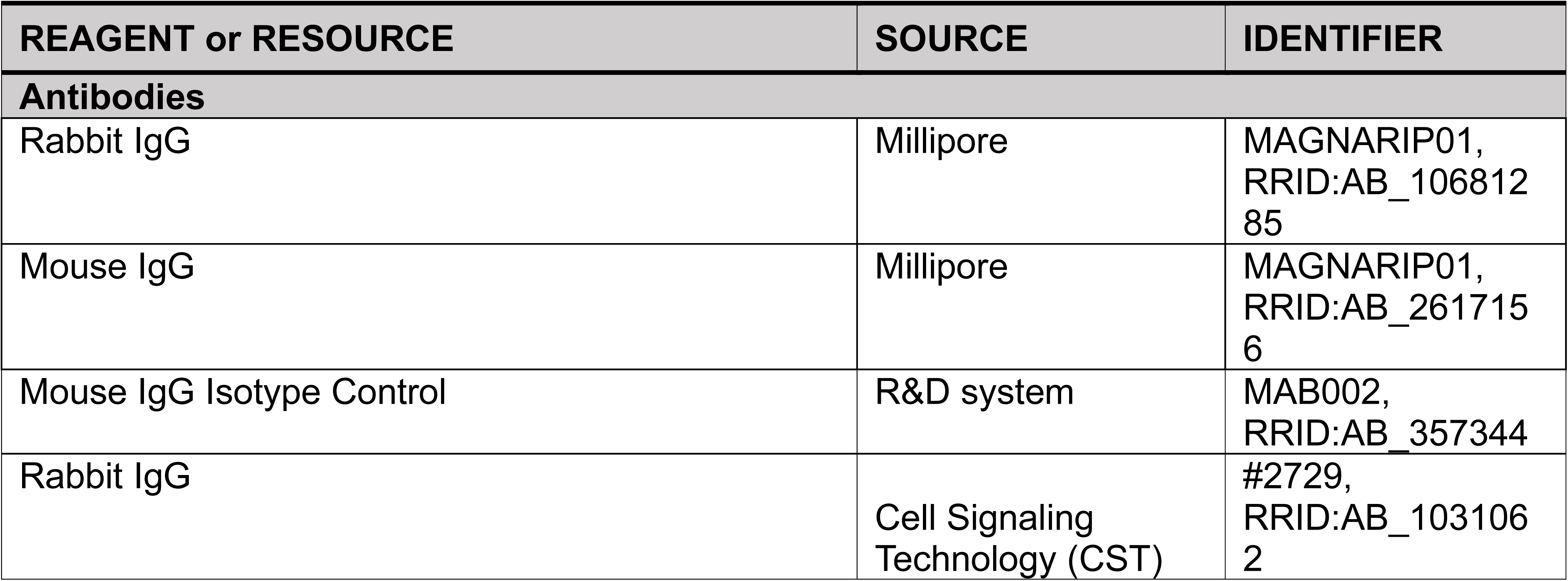
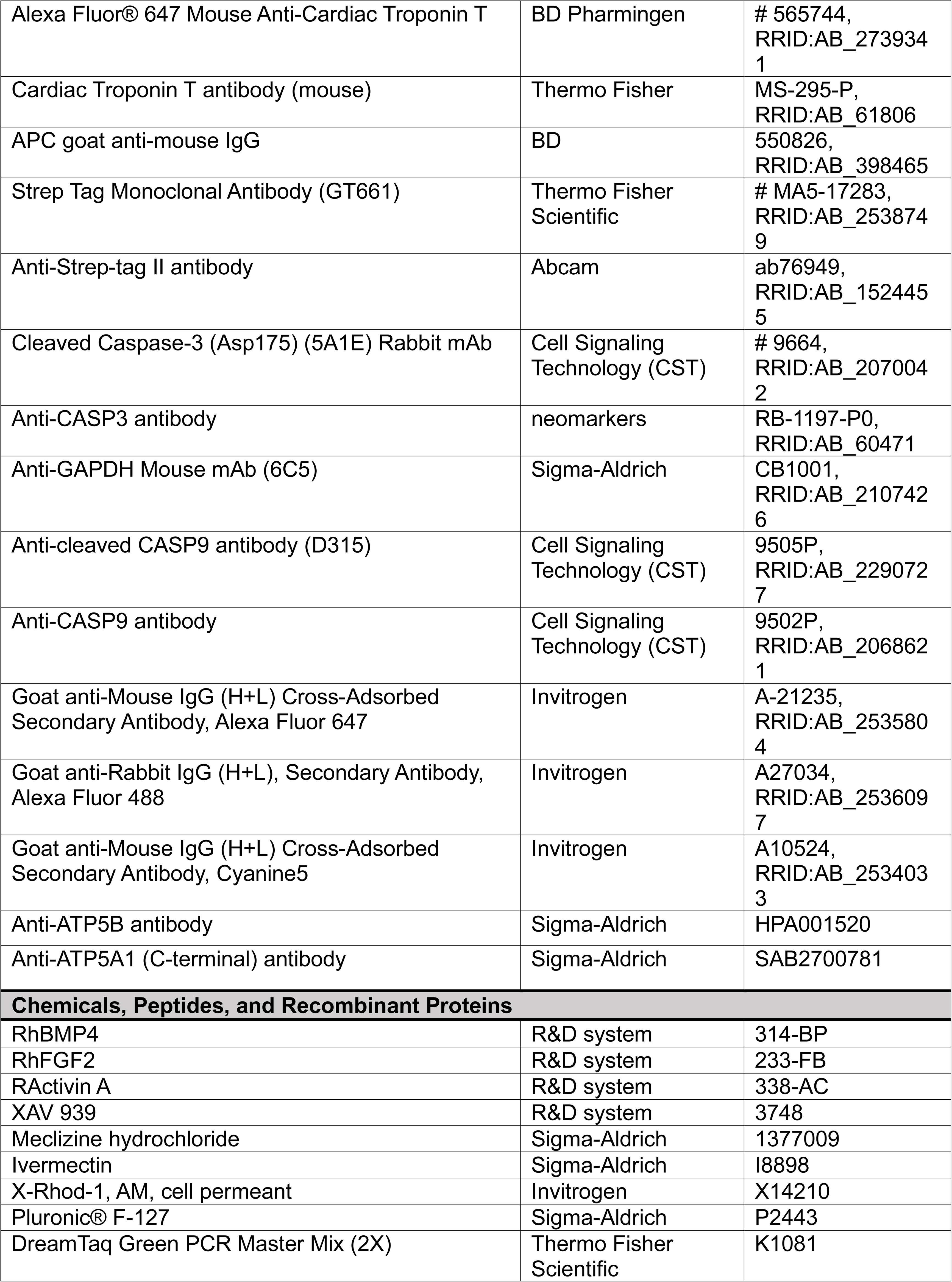
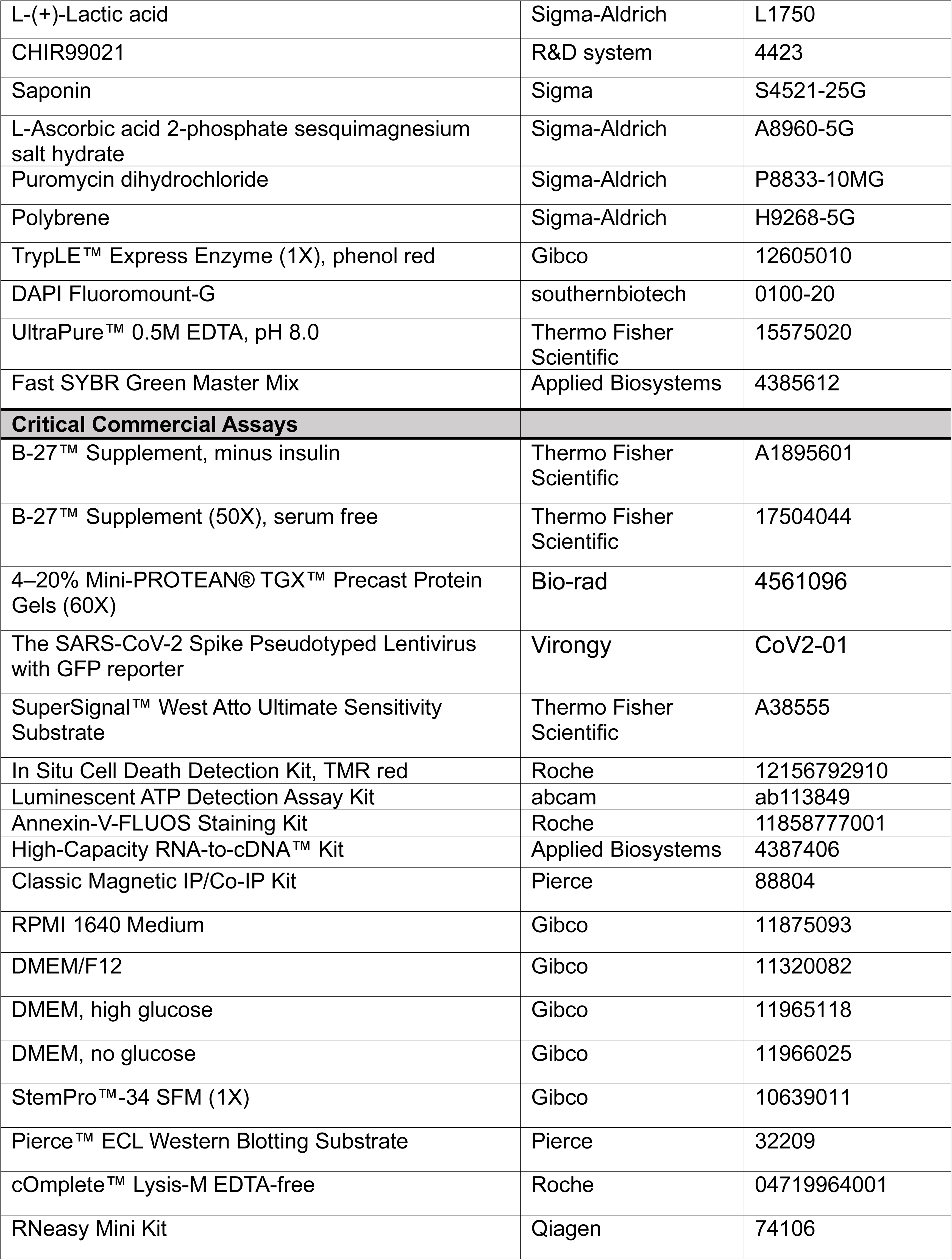
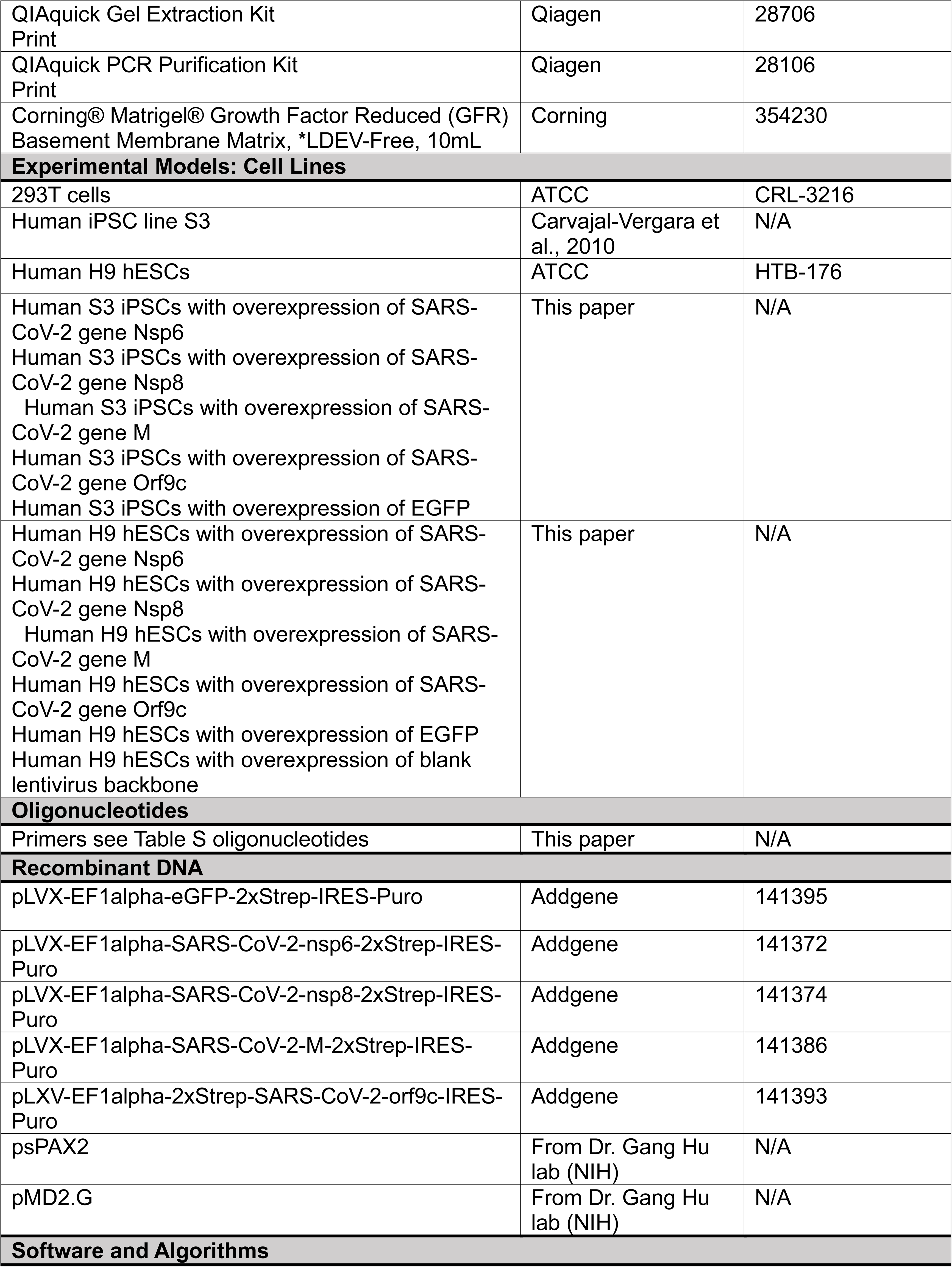
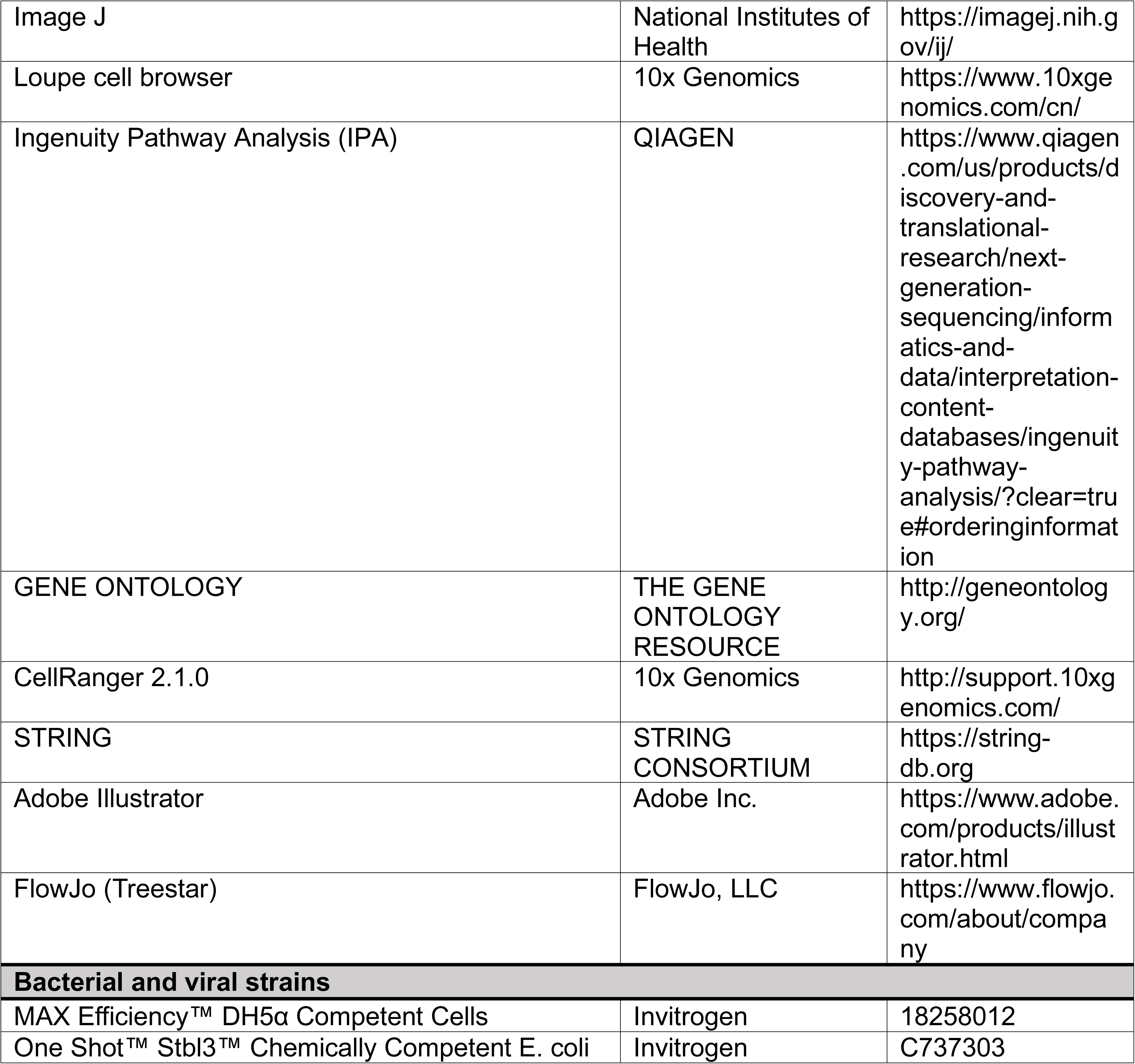

| REAGENT or RESOURCE | SOURCE | IDENTIFIER |
| --- | --- | --- |
| **Antibodies** |  |  |
| Rabbit IgG | Millipore | MAGNARIP01, RRID:AB_10681285 |
| Mouse IgG | Millipore | MAGNARIP01, RRID:AB_2617156 |
| Mouse IgG Isotype Control | R&D system | MAB002, RRID:AB_357344 |
| Rabbit IgG | Cell Signaling Technology (CST) | #2729, RRID:AB_1031062 |
| Alexa Fluor® 647 Mouse Anti-Cardiac Troponin T | BD Pharmingen | # 565744,<br>RRID:AB_273934<br>1 |
| Cardiac Troponin T antibody (mouse) | Thermo Fisher | MS-295-P,<br>RRID:AB_61806 |
| APC goat anti-mouse IgG | BD | 550826,<br>RRID:AB_398465 |
| Strep Tag Monoclonal Antibody (GT661) | Thermo Fisher Scientific | # MA5-17283,<br>RRID:AB_253874<br>9 |
| Anti-Strep-tag II antibody | Abcam | ab76949,<br>RRID:AB_152445<br>5 |
| Cleaved Caspase-3 (Asp175) (5A1E) Rabbit mAb | Cell Signaling Technology (CST) | # 9664,<br>RRID:AB_207004<br>2 |
| Anti-CASP3 antibody | neomarkers | RB-1197-P0,<br>RRID:AB_60471 |
| Anti-GAPDH Mouse mAb (6C5) | Sigma-Aldrich | CB1001,<br>RRID:AB_210742<br>6 |
| Anti-cleaved CASP9 antibody (D315) | Cell Signaling Technology (CST) | 9505P,<br>RRID:AB_229072<br>7 |
| Anti-CASP9 antibody | Cell Signaling Technology (CST) | 9502P,<br>RRID:AB_206862<br>1 |
| Goat anti-Mouse IgG (H+L) Cross-Adsorbed Secondary Antibody, Alexa Fluor 647 | Invitrogen | A-21235,<br>RRID:AB_253580<br>4 |
| Goat anti-Rabbit IgG (H+L), Secondary Antibody, Alexa Fluor 488 | Invitrogen | A27034,<br>RRID:AB_253609<br>7 |
| Goat anti-Mouse IgG (H+L) Cross-Adsorbed Secondary Antibody, Cyanine5 | Invitrogen | A10524,<br>RRID:AB_253403<br>3 |
| Anti-ATP5B antibody | Sigma-Aldrich | HPA001520 |
| Anti-ATP5A1 (C-terminal) antibody | Sigma-Aldrich | SAB2700781 |
| **Chemicals, Peptides, and Recombinant Proteins** |  |  |
| RhBMP4 | R&D system | 314-BP |
| RhFGF2 | R&D system | 233-FB |
| RActivin A | R&D system | 338-AC |
| XAV 939 | R&D system | 3748 |
| Meclizine hydrochloride | Sigma-Aldrich | 1377009 |
| Ivermectin | Sigma-Aldrich | I8898 |
| X-Rhod-1, AM, cell permeant | Invitrogen | X14210 |
| Pluronic® F-127 | Sigma-Aldrich | P2443 |
| DreamTaq Green PCR Master Mix (2X) | Thermo Fisher Scientific | K1081 |
| L-(+)-Lactic acid | Sigma-Aldrich | L1750 |
| CHIR99021 | R&D system | 4423 |
| Saponin | Sigma | S4521-25G |
| L-Ascorbic acid 2-phosphate sesquimagnesium salt hydrate | Sigma-Aldrich | A8960-5G |
| Puromycin dihydrochloride | Sigma-Aldrich | P8833-10MG |
| Polybrene | Sigma-Aldrich | H9268-5G |
| TrypLE™ Express Enzyme (1X), phenol red | Gibco | 12605010 |
| DAPI Fluoromount-G | southernbiotech | 0100-20 |
| UltraPure™ 0.5M EDTA, pH 8.0 | Thermo Fisher Scientific | 15575020 |
| Fast SYBR Green Master Mix | Applied Biosystems | 4385612 |
| **Critical Commercial Assays** |  |  |
| B-27™ Supplement, minus insulin | Thermo Fisher Scientific | A1895601 |
| B-27™ Supplement (50X), serum free | Thermo Fisher Scientific | 17504044 |
| 4–20% Mini-PROTEAN® TGX™ Precast Protein Gels (60X) | Bio-rad | 4561096 |
| The SARS-CoV-2 Spike Pseudotyped Lentivirus with GFP reporter | Virongy | CoV2-01 |
| SuperSignal™ West Atto Ultimate Sensitivity Substrate | Thermo Fisher Scientific | A38555 |
| In Situ Cell Death Detection Kit, TMR red | Roche | 12156792910 |
| Luminescent ATP Detection Assay Kit | abcam | ab113849 |
| Annexin-V-FLUOS Staining Kit | Roche | 11858777001 |
| High-Capacity RNA-to-cDNA™ Kit | Applied Biosystems | 4387406 |
| Classic Magnetic IP/Co-IP Kit | Pierce | 88804 |
| RPMI 1640 Medium | Gibco | 11875093 |
| DMEM/F12 | Gibco | 11320082 |
| DMEM, high glucose | Gibco | 11965118 |
| DMEM, no glucose | Gibco | 11966025 |
| StemPro™-34 SFM (1X) | Gibco | 10639011 |
| Pierce™ ECL Western Blotting Substrate | Pierce | 32209 |
| cOmplete™ Lysis-M EDTA-free | Roche | 04719964001 |
| RNeasy Mini Kit | Qiagen | 74106 |
| QIAquick Gel Extraction Kit<br>Print | Qiagen | 28706 |
| QIAquick PCR Purification Kit<br>Print | Qiagen | 28106 |
| Corning® Matrigel® Growth Factor Reduced (GFR)<br>Basement Membrane Matrix, *LDEV-Free, 10mL | Corning | 354230 |
| **Experimental Models: Cell Lines** |  |  |
| 293T cells | ATCC | CRL-3216 |
| Human iPSC line S3 | Carvajal-Vergara et al., 2010 | N/A |
| Human H9 hESCs | ATCC | HTB-176 |
| Human S3 iPSCs with overexpression of SARS-CoV-2 gene Nsp6<br>Human S3 iPSCs with overexpression of SARS-CoV-2 gene Nsp8<br>Human S3 iPSCs with overexpression of SARS-CoV-2 gene M<br>Human S3 iPSCs with overexpression of SARS-CoV-2 gene Orf9c<br>Human S3 iPSCs with overexpression of EGFP | This paper | N/A |
| Human H9 hESCs with overexpression of SARS-CoV-2 gene Nsp6<br>Human H9 hESCs with overexpression of SARS-CoV-2 gene Nsp8<br>Human H9 hESCs with overexpression of SARS-CoV-2 gene M<br>Human H9 hESCs with overexpression of SARS-CoV-2 gene Orf9c<br>Human H9 hESCs with overexpression of EGFP<br>Human H9 hESCs with overexpression of blank lentivirus backbone | This paper | N/A |
| **Oligonucleotides** |  |  |
| Primers see Table S oligonucleotides | This paper | N/A |
| **Recombinant DNA** |  |  |
| pLVX-EF1alpha-eGFP-2xStrep-IRES-Puro | Addgene | 141395 |
| pLVX-EF1alpha-SARS-CoV-2-nsp6-2xStrep-IRES-Puro | Addgene | 141372 |
| pLVX-EF1alpha-SARS-CoV-2-nsp8-2xStrep-IRES-Puro | Addgene | 141374 |
| pLVX-EF1alpha-SARS-CoV-2-M-2xStrep-IRES-Puro | Addgene | 141386 |
| pLVX-EF1alpha-2xStrep-SARS-CoV-2-orf9c-IRES-Puro | Addgene | 141393 |
| psPAX2 | From Dr. Gang Hu lab (NIH) | N/A |
| pMD2.G | From Dr. Gang Hu lab (NIH) | N/A |
| **Software and Algorithms** |  |  |
| Image J | National Institutes of Health | https://imagej.nih.gov/ij/ |
| Loupe cell browser | 10x Genomics | https://www.10xgenomics.com/cn/ |
| Ingenuity Pathway Analysis (IPA) | QIAGEN | https://www.qiagen.com/us/products/discovery-and-translational-research/next-generation-sequencing/informatics-and-data/interpretation-content-databases/ingenuity-pathway-analysis/?clear=true#orderinginformation |
| GENE ONTOLOGY | THE GENE ONTOLOGY RESOURCE | http://geneontology.org/ |
| CellRanger 2.1.0 | 10x Genomics | http://support.10xgenomics.com/ |
| STRING | STRING CONSORTIUM | https://string-db.org |
| Adobe Illustrator | Adobe Inc. | https://www.adobe.com/products/illustrator.html |
| FlowJo (Treestar) | FlowJo, LLC | https://www.flowjo.com/about/company |
| **Bacterial and viral strains** |  |  |
| MAX Efficiency™ DH5α Competent Cells | Invitrogen | 18258012 |
| One Shot™ Stbl3™ Chemically Competent E. coli | Invitrogen | C737303 |

